# Optogenetic strategies for optimizing the performance of biosensors of membrane phospholipids in live cells

**DOI:** 10.1101/2023.08.03.551799

**Authors:** Yuanfa Yao, Xiayan Lou, Luhong Jin, Weiyun Sun, Jingfang Liu, Yunyue Chen, Sunying Cheng, Tengjiao Zhao, Shuwei Ke, Luhao Zhang, Yingke Xu, Lian He, Hanbing Li

**Author notes:** These authors contributed equally: Yuanfa Yao, Xiayan Lou. Correspondence should be addressed to **Drs. Hanbing Li**, **Drs. Lian He**, **Prof. Yinek Xu**, **Address for Dr. Hanbing Li (lead):** Institute of Pharmacology, College of Pharmaceutical Science, 18 Chaowang Road, Zhejiang University of Technology, Hangzhou, 310014, China; Address for Dr. Lian He: Department of Pharmacology, School of Medicine, 1088 Xueyuan Avenue, Southern University of Science and Technology, Shenzhen, 518055, China.; Address for Prof. Yingke Xu: Department of Biomedical Engineering, 38 Zheda Road, Yuquan Campus, Zhejiang University, Hangzhou, 310027, China.

## Abstract

High-performance biosensors play a crucial role in elucidating the intricate spatiotemporal regulatory roles and dynamics of membrane lipids. However, enhancing their sensitivity and substrate-detecting capabilities remains a significant challenge. Here, we presented optogenetic-based strategies to optimize phospholipid biosensors. These strategies involved pre-sequestering unbound probes in the cell nucleus to minimize background signals in the cytoplasm. These stored probes could be released from the nucleus in response to blue light according to experimental requirements. Furthermore, we employed optically-controlled phase separation to generate punctate probes that amplified signals and facilitated the visualization of phospholipids in cells. These improved phospholipid biosensors hold great potential for enhancing our understanding of the spatiotemporal dynamics and regulatory roles of membrane lipids in live cells and this methodological insights might be valuable for developing other high-performance biosensors.

## 1. Introduction

The dynamic metabolism of membrane lipids is a rapid and intricate process that demands high-performance probes for real-time monitoring within living cells. Biosensors play a crucial role in unraveling the spatiotemporal regulatory functions of membrane lipids and have been widely employed to track their fluctuations within cellular environments. A primary optimization strategy for enhancing these biosensors involves selecting sensitive phospholipid-binding motifs from a variety of proteins of interest (POI). However, under certain conditions, such as detecting low-abundance phospholipids, the unbound fraction resulting from biosensor overexpression within cells can exert varying degrees of influence on detection performance. In contrast to those efforts aimed at enhancing biosensor sensitivity, there has been relatively little exploration into reducing background signals from the unbound biosensors to improve their signal-to-noise ratio (SNR) or amplifying these signals during the imaging process.

What methods can be employed to more effectively eliminate unbound biosensors and achieve a clearer subcellular visualization of membrane lipid distribution? A more effective strategy is to segregate dissociated probes to the specific region and minimize their impact on the SNR in imaging process. For example, secluding the unbound probes to the nucleus through optogenetic tools, such as LOV2-based nucleocytoplasmic transport system^1,2^. Optogenetics has found extensive application in manipulating the spatiotemporal localization^3-6^ and conformational dynamics of POIs^7^, and regulating gene transcription^8-10^, modulating ion channel states^11^, and facilitating the intracellular movement of organelles^12,13^ in numerous studies. Its high temporal-spatial resolution and reversibility empowers us to manipulate complex cellular processes, leading to a deeper comprehension of the mechanical principles governing both physiological and pathological activities. With the regard to amplify the signal of biosensors, the SunTag system and similar tools has been extensively used to amplify the single-molecular signal through incorporation of a dozen of tandem short peptides, such as GCN4^14-16^, HA^17^, and gp41 peptide^18^. These peptides are then recognized by a multitude of fluorescent antibodies, resulting in the formation of punctum-like structures that are notably brighter than the background, thus amplifying the signal. Importantly, this process requires to control the expression levels of short peptides by doxycycline (DOX) induction. Recently, puncta induced by Homo-Oligomeric Tags (HOTags)-mediated phase separation have also been utilized to report the cellular activities^19-21^. The formation of these puncta is dependent on the interactions of fused proteins with HOTags. The brightness of them is comparative to that of the SunTag system.

It remains unclear whether the aforementioned advanced techniques can enhance the performance of phospholipids biosensors. In pursuit of addressing this question, our study employed cpLOV2-mediated nucleocytoplasmic transport to modify phospholipid biosensors. This mechanism sequesters unbound biosensors within the cell nucleus under dark conditions via NLS-mediated nuclear entry, leading to a significant reduction in background noise and an improvement in SNR of the biosensors. Upon subsequent exposure to light, these stored probes were released from the nucleus, ready to bind to newly-synthesized phospholipids in response to agonists. We applied these strategies to enhance the performance of phosphatidic acid (PA) biosensor. Moreover, to further improve biosensors, we used CRY2/CIBN system to optically trigger the phase separation of HOTag3 and HOTag6^19-21^. This novel approach not only amplified the signal of the phosphatidylinositol-4-phosphate (PI4P) and PA biosensors but also rapidly assembled a cluster of biosensors with multiple copies of phospholipid-binding motifs, enhancing their sensitivity to respective phospholipids. This differs from the construction of repetitive sequences of the binding motif within a single biosensor. The punctum-like structure of the biosensors facilitated the visualization of intracellular phospholipids and allowed us to readily observe low-abundance phospholipids on cellular organelles and detect minor concentration changes. Collectively, these refined biosensors may serve as useful tools for investigating the biological functions of PI4P and PA. The methodological improvements presented in this study can also serve as valuable references for the design of high-performance biosensors.

## 2. Results and Discussion

### 2.1 Designing optogenetics-based biosensors with circularly permuted LOV2 (cpLOV2) to minimize interference from unbound biosensors

In previous reports and reviews, we highlighted the significant roles of PLDs and PA in cells^22-24^, especially in cancer cells. However, to gain a comprehensive understanding of their spatiotemporal regulatory roles, we need to monitor their real-time activity and thus advanced detection methods are indispensable. Baskin’s group developed a sophisticated chemical detection platform based on click chemistry for labeling PA and tracking PLD activities^25^. This platform facilitated the investigation of localization-dependent regulatory roles of PLDs^26^ and identification of cells with heightened PLD activities^27^. Apart from this technique, detection of other membrane lipids predominantly relies on biosensors in live cells, employing specific motifs from phospholipid-binding proteins. However, the presence of unbound biosensors can significantly impact the imaging process, especially with low-affinity probes and detecting low-abundance substrates.

To minimize the their impacts, we integrated an optogenetic element to sequester unbound molecules into the nucleus. To do this, we first created a cpLOV2-mCherry (mCh)-NLS construct and added different nuclear export sequence 1 (NES1) at the N-terminal Jα helix (**Fig. S1A**). Under the dark condition, the majority of these proteins were localized within the cell nucleus ascribe to the presence of NLS (**Fig. S1B**). After exposure to blue light, the N-terminal Jα helix was supposed to unfold and expose the fused NES to mediate the nuclear export. Two out of the seven tested NES sequences exhibited the expected nuclear export and the NES (DEAAKELAGLDL) demonstrated the highest effectiveness (**Fig. S1A-B**). Subsequently, we conducted a comparative analysis of the PA binding capabilities of different PA binding domains (PABDs). Notably, the PABD from yeast overproduction of inositol 1 gene product (Opi1p) and human Raf1 did not perform as anticipated in detecting PA levels at the plasma membrane (**Fig. S1C-D**). Consequently, we selected the PABD from the PA biosensor PASS. This wildly-used probe were developed by Du’s group and used the PABD (51-91) from SPO20 for PA-binding^28^. They inserted a NES to counteract to the nuclear localization of PABD, thus promoting increased expression of the biosensor in the cytosol. Owing to the potential interference of this NES with the action of NLS that intended to seclude the unbound biosensors into the nucleus, we mutated two key hydrophobic residues (leucine and isoleucine) into alanine to deactivate this NES (NES2). The optimized cpLOV2 was added to the N-terminal of the PASS biosensor and we introduced two or more tandem copies of PABD to increase the PA-binding ability (**Fig. 1A**). This optogenetics-based PA biosensor was subsequently renamed as **Opto**genetics **PA S**uper **S**ensor based on **c**pLOV2 (Opto-PASSc) (**Fig. 1A**).

**Figure 1.**
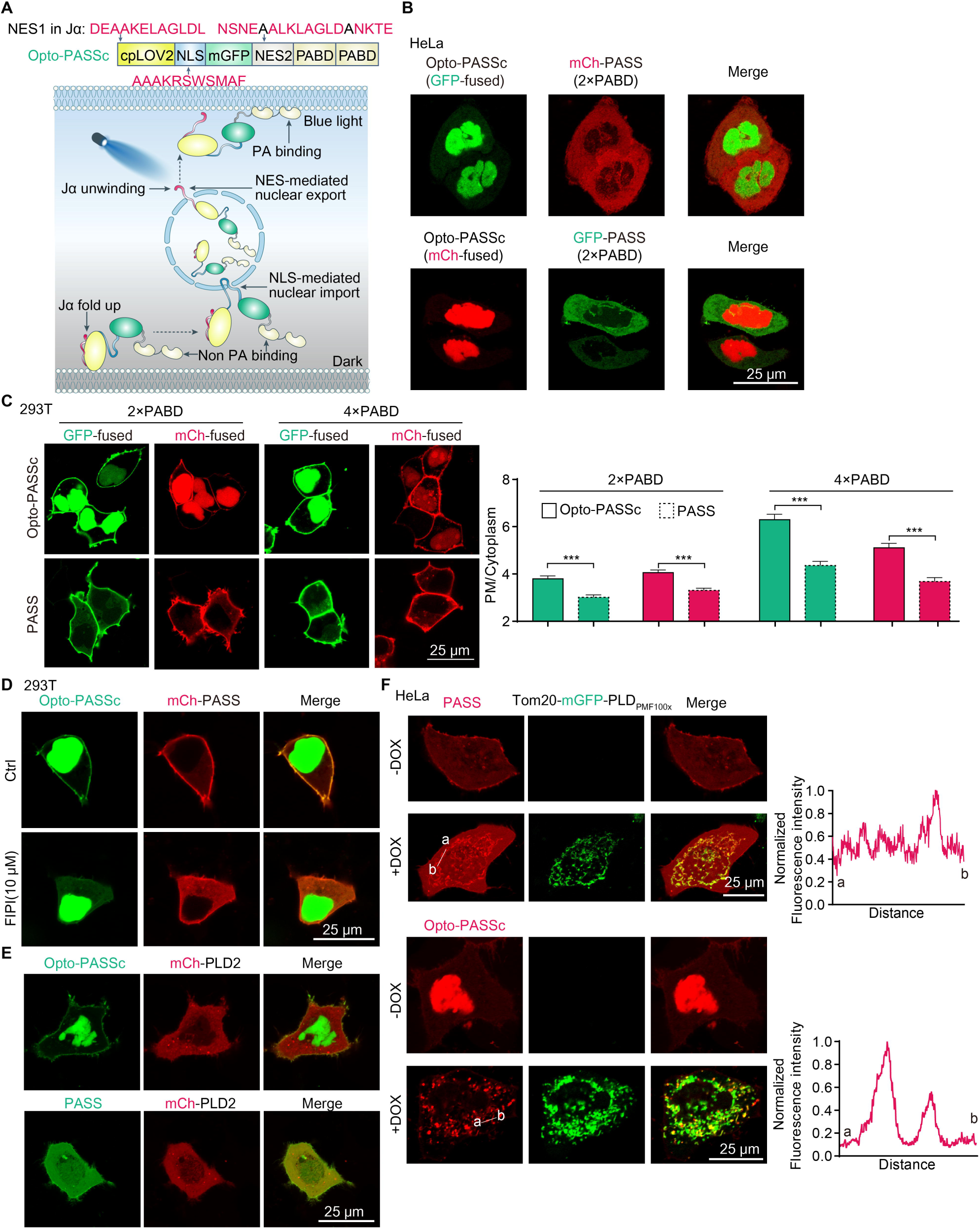
Design of optogenetics-based phospholipid biosensor with circularly permuted LOV2 (cpLOV2). **A**, a schematic diagram of the cpLOV-based strategy for improvements of membrane phospholipids biosensors, NES1 was inserted in the Jα helix of the cpLOV2 element and exposed upon blue light stimulation; two residues (Leucine and Isoleucine) of the NES2 was mutated into Alanine and colored in black. **B**, the intracellular localization of Opto-PASSc (fused with Emerald (GFP) or mCherry (mCh)) with PASS biosensor (two copies of PABDs) in HeLa cells. **C**, representative images of Opto-PASSc and PASS with two or four copies of PABDs and the fluorescence intensity ratio of plasma membrane (PM) to the cytosol was used to assess their PA-detecting ability in 293T cells (n ≥ 45 cells). Opto-PASSc and PASS were used to reflect PA levels at the PM after four-hour treatment of PLDs inhibitor FIPI (10 μM) in 293T cells (**D**) and overexpression of PLD2 in HeLa cells (**E**). **F**, Opto-PASSc and PASS in detecting PA on the mitochondria through a tetracycline-inducible expression of highly-active PLD_PMF100×_, doxycycline (2 μg/ml) incubation time was five hours; the normalized fluorescence intensity of the white line (ab) was analyzed to assess the signal-to-noise of these two biosensors.

The mutation of NES2 further increased the NLS-mediated nuclear localization of Opto-PASSc (**Fig. S2A-B**) and these probes predominantly localized to the nucleus (**Fig. 1B**), suggesting a low basal levels of PA. Additionally, they also exhibited colocalization with the referenced PA probes RFP-PASS and GFP-PASS in HeLa and 293T cells (**Fig. S2C-D**). In the case of Opto-PASSc with two copies of PABDs, the different fused fluorescent proteins had limited impact on the PA-detecting ability at the plasma membrane (**Fig. 1C** and **Fig. S2E**), while GFP-labelled Opto-PASSc outperformed the mCh-fused biosensors when increasing the number of the PABD copies. Moreover, Opto-PASSc demonstrated a lower background signal in the cytosol, and the fluorescence intensity ratio of plasma membrane (PM) to cytosol was higher in cells expressing Opto-PASSc compared to PASS. This partially indicates an enhancement in the PA-detecting performance of Opto-PASSc. Additionally, we also generated an Opto-PASSc variant with two mutated PABDs that is incapable of binding PA. The results demonstrated that this construct exhibited reduced localization at the PM compared to Opto-PASSc with wild-type PABDs (**Fig. S1E**), partially suggesting its specificity for PA binding.

Both Opto-PASSc and PASS exhibited high basal levels of PA at the plasma membrane in 293T cells. As PLD1 and PLD2 are main enzymes that biosynthesize PA by hydrolysis of phosphatidylcholine (PC), we used FIPI, a dual inhibitor of PLD1 and PLD2, to abolish their action on PA production. Both these two biosensors reflected PA reduction at the PM after a four-hour FIPI treatment (**Fig. 1D**). Additionally, we observed that Opto-PASSc characterized the increase in PA at the PM more clearly after overexpression of PLD2 compared to the PASS biosensor (**Fig. 1E**).

Given that the basal activities of endogenous PLDs were typically low, their activation requires additional stimulation^23,24,29^, such as G-protein-coupled receptor agonists and PKC activation. To assess the PA-detecting performance of our enhanced biosensor in the cytosol, we generated a tetracycline-inducible construct of PLD with elevated basal activity (referred to here as PLD_PMF100×_). This variant was called SuperPLD and was created by Baskin’s group^30^. Its basal activity is a hundredfold higher than that of the wild-type PLD_PMF_. After a five-hour induction with DOX, we could clearly observe that the Tom20-fused PLD_PMF100×_ is located on the mitochondria in contrast to the untreated group. This resulted in the production of PA on the mitochondria, which was detected by both the Opto-PASSc and PASS biosensors (**Fig. 1F**). We analyzed the SNR of these biosensors, respectively and the Opto-PASSc displayed a higher SNR, which is consisted with aforementioned results of PA-detecting at the PM (**Fig. 1F**). Moreover, the signal of Opto-PASSc in the nucleus was distinctly reduced in response to increasing PLD activities in the cytosol (**Fig. 1F**). This indicates that Opto-PASSc exhibited a preference for binding to PA rather than entering the nucleus and sequestering them in the nucleus by NLS has limited impact on their ability to bind to PA under conditions where PA levels are extremely elevated in the cytosol.

### 2.2 Light-induced nuclear export of Opto-PASSc enhanced the PA-detecting performance

We first investigated whether blue light induced translocation of Opto-PASSc into the cytosol. The N-terminal Jα helix fused with the NES was expected to unwind in response to blue light stimulation, exposing the NES and mediating the nuclear export of biosensors. Consequently, more probes were transported into the cytosol for binding to PA. By analyzing the fluorescence intensity ratio of the cytoplasm to the nucleus, we observed an increase in this ratio in response to consecutive blue light pulse stimulation (**Fig. 2A**). Compared to the construct lacking fusion of NES into the Jα helix of cpLOV2, the signals in the nucleus decreased after blue light illumination (**Fig. 2B**).

**Figure 2.**
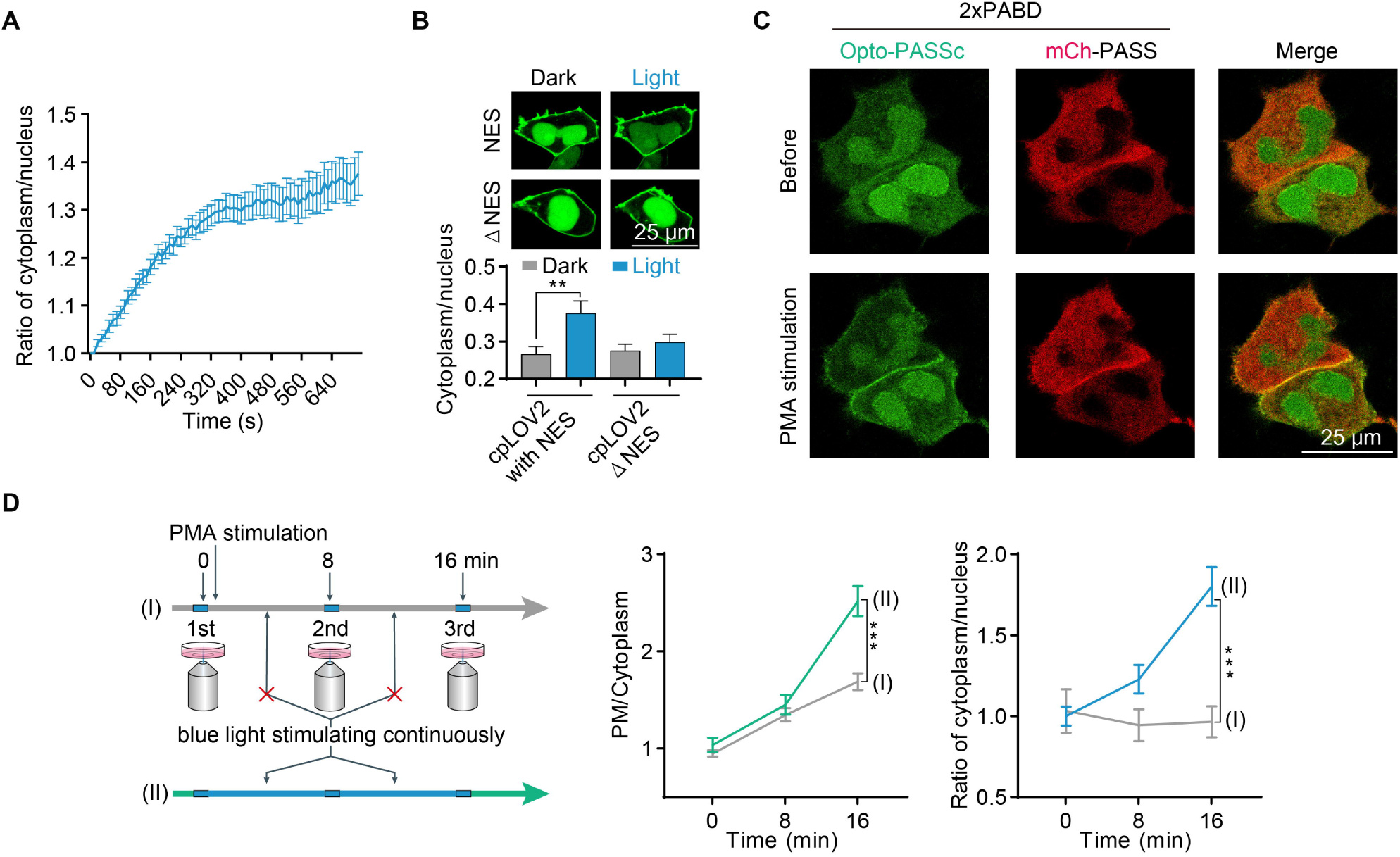
Blue light stimulation induced the nuclear export and enhanced the PA-detecting ability of Opto-PASSc biosensor. **A**, the fluorescence intensity ratio of the cytosol to nucleus was analyzed to indicate the nuclear export of Opto-PASSc in responsive to blue light stimulation (2% rated power of 488 nm laser, pulse interval was 10 second), the data was normalized to the initial value (n ≥ 7 cells). **B**, the nuclear export of Opto-PASSc with or without NES (ΔNES) in the Jα helix of the cpLOV2 in responsive to blue light stimulation in 293T cells (n ≥ 23 cells). **C**, Opto-PASSc and PASS (2×PABD) in detecting PA changes before and after indirect PLD activator PMA (5 μM) treatment for 16 minutes. **D**, the impact of blue light stimulation on the PA-detecting performance of Opto-PASSc in response to PMA (5 μM)-induced PA production, the condition (I): only three images were captured at indicated time and the interval was dark treatment; the condition (II): blue light stimulation at the capture interval, PMA was added after finishing capture of the first image, the data was normalized to the mean of initial value (n ≥ 20 cells).

To evaluate whether the light-induced export of Opto-PASSc could enhance its performance in reflecting PA production, we employed an indirect activator, phorbol 12-myristate 13-acetate (PMA) (a PKC activator), to instantly increase PA levels. We observed PA levels at the PM was elevated after incubation of PMA (**Fig. 2C**). Considering the short stimulation time of blue light during the acquisition of GFP channel signals, which is insufficient to induce a significant export of the probe from the nucleus, we compared the effects of two treatments: the dark condition and consecutive blue light stimulation at capture intervals, on the PA-detecting capability of Opto-PASSc in cells responding to PMA treatment. Our findings revealed that consecutive blue light stimulation at capture intervals reduced the levels of Opto-PASSc in the nucleus compared to the dark condition (**Fig. 2D** and **Fig. S2F**). Moreover, the fluorescence intensity ratio (PM to the cytoplasm) was higher under this condition (**Fig. 2D** and **Fig. S2F**), suggesting that light illumination enhanced the detection performance of Opto-PASSc in monitoring instantaneous PA production at the PM.

Collectively, Opto-PASSc consistently demonstrates a low background signal by pre-sequestering unbound probes in the cell nucleus. This feature provides an accurate reflection of PA levels in the cytosol under basal conditions, a capability not present in the traditional PA biosensor PASS. Importantly, this improvement does not compromise the PA-detecting ability of Opto-PASSc, as it still preferentially binds to PA rather than extensively entering the nucleus in response to cytoplasmic PA increases. Additionally, the nucleocytoplasmic transport of Opto-PASSc can be controlled by light according to experimental requirements, indicating greater flexibility in imaging PA in living cells under various experimental conditions.

### 2.3 Development of optically-controlled phase separation to amplify the signal of biosensors

Biosensors with high-resolution have the potential to significantly advance our understanding of the specific mechanistic roles of membrane lipids in live cells. In addition to utilizing super-resolution microscopy equipment, amplifying the fluorescence signal of biosensors offers an alternative method to visualize the dynamics of probes inside cells. Therefore, we exploited the interaction-sensitive phase separation of the HOTag3/HOTag6 system to generate punctum-like structures, thereby amplifying the fluorescence signal of biosensors in cells^19^. In this system, HOTag3 can assemble into a hexamer and HOTag6 forms a tetramer. The phase separation would be triggered by the interaction of their fused fragments. By virtue of the light-responsive binding of the optogenetic tool CRY2/CIBN, the phase separation process of HOTag3/HOTag6 system became controllable via light stimulation (**Fig. 3A**). Then we added a phospholipid-binding motif, such as PABD and P4M domain from SidM, to the CIBN fragment for detecting corresponding phospholipid. In this way, this improved biosensor system generates fluorescent puncta in response to light. The punctum-like probe can bind to intracellular PI4P and it enables us to visually observed the spatial distribution and precise positional information of PI4P more intuitively, which consequently improves its detection performance in comparison to the sole expression of the fluorescin-fused phospholipid-binding domain (**Fig. 3A**). These improved biosensors were called **Opto**genetic **PI4P s**ensor or **PA s**uper **s**ensor based on **p**hase **s**eparation (Opto-PI4PSps or Opto-PASSps) (**Fig. 3A**).

**Figure 3.**
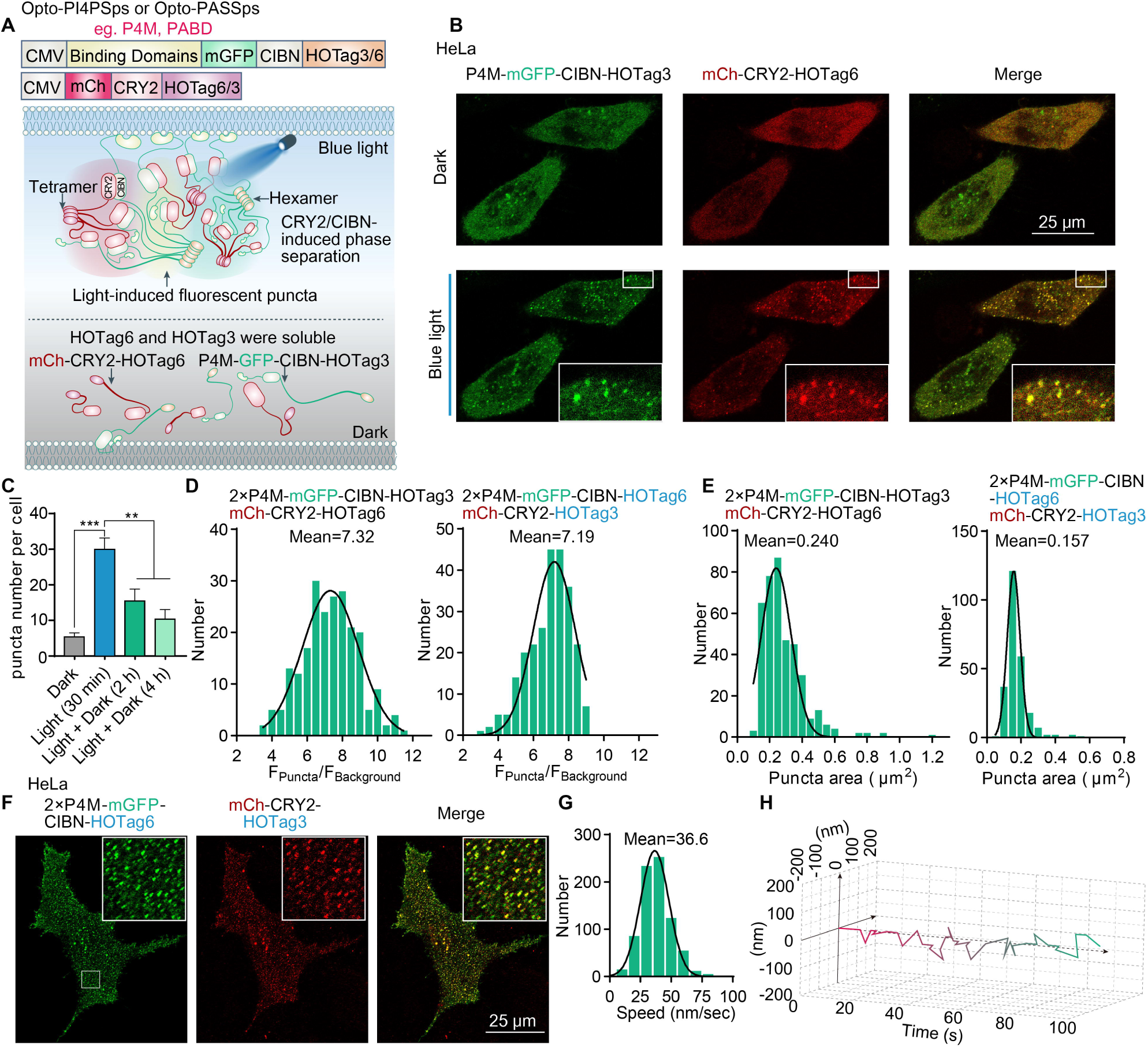
amplification of biosensors signal based on the optically-controlled phase separation. **A**, a schematic diagram illustrating optically-controlled phase separation for optimizing membrane phospholipid biosensors. **B**, the light-induced binding of CRY2/CIBN triggered the phase separation via HOTag3/HOTag6 in HeLa cells (blue light stimulation: 2% rated power of 488 nm laser for 10 min and the pulse interval was 5 seconds). **C**, the phase separation puncta slowly disaggregated under dark condition (n ≥ 40 cells). **D**, the histogram reflected the brightness distribution of phase separation spots by measuring the fluorescence intensity ratio of puncta to the background in HeLa cells with co-expression of indicated plasmids and **E**, their the planar dimensions of phase separation puncta (n ≥ 200 puncta). **F**, the representative images of phase separation puncta depicted the distribution of PI4P on the plasma membrane after light stimulation in HeLa cells and **G**, the speed histogram reflects the movement of these puncta (n ≥ 700), and **H**, the trajectory plot of a representative phase separation punctum.

The results were consistent with our expectations. We observed a 10-minute light pulse stimulation induced both green and red puncta in cells (**Fig. 3B**), even after we exchanged the HOTag3 and HOTag6 to the corresponding fused proteins (**Fig. S3A**). To further assess the reversibility of light-induced phase separation, we incubated the stimulated cells in the absence of light for several hours. Our observations revealed that a portion of the phase separation puncta diffused without the presence of light stimulation (**Fig. S3B** and **Fig. 3C**). Given that the disassociation time of CRY2 and CIBN is approximately 10-25 minutes^31^, we speculated that the diffusion kinetics of phase separation puncta are primarily determined by the disaggregation of HOTag3 and HOTag6. In addition, we also excluded these puncta were self-oligomerization of CRY2 (**Fig. S3C**). Our aforementioned result indicated addition of an extra copy of the P4M domain extremely increased its affinity (**Fig. S3D**). Consequently, we fused two copies of the P4M domain with CIBN-HOTag3 or CIBN-HOTag6. The fluorescence intensity of these puncta was approximately 7.32-fold higher than the background signal in the GFP channel (**Fig. 3D**). Their planar size was approximately 0.240 μm^2^ (**Fig. 3E**). Intriguingly, upon exchanging the fusion of HOTag3 and HOTag6 with their respective optogenetic counterparts (as indicated in the text), there was a modest reduction in both the brightness and size of these puncta (**Fig. 3D-E**).

Next, we investigated whether this technological refinement enhanced the detection performance of the PI4P biosensor at the PM. Our analysis indicated that the majority of these spots at the PM had velocities approximately at 36.6 nm/s (**Fig. 3F-H**), a rate significantly lower than the translocation speed of kinesin (635-1,290 nm/s) as reported by reference^14^. This data suggests that the majority of the spots at the PM exhibit small-scale movements under the basal state and our probe potentially offer partial insights into the dynamics of PI4P at the PM. In addtition, each punctum incorporated a dozen of phospholipid-binding motifs, which indicated our improved biosensors had higher affinity to the targeted phospholipids. However, the subsequent limitation is the challenge in determining the exact number of phospholipid molecules bound to a single punctum. It might be regulated by selecting appropriate HOTags to fuse with phospholipid-binding motifs. For instance, as HOTag6 forms a tetramer, the resulting cluster may have fewer probe heads available for phospholipid binding compared to the design utilizing the hexameric HOTag3.

### 2.4 Opto-PI4PSps and Opto-PASSps biosensors allowed for the visualization of intracellular phospholipids

Both Opto-PI4PSps and Opto-PASSps biosensors colocalized with the non-optimized traditional biosensor (**Fig. 4A-B**), indicating that phase separation-mediated signal amplification does not hinder the binding of biosensors to their respective phospholipids. These two biosensors were able to visually indicate PI4P and PA at the PM (**Fig. 4A-B**). For example, through the detection by Opto-PI4PSp, we can visually observed the reduction of PI4P levels at the PM after treatment of a inhibitor phenylarsine oxide (PAO) to blockage the production of PI4P (**Fig. S3E**). With the aid of this optimized probe, we clearly observed that certain punctate probes were co-localized with lysosomes and moved synchronously with them (**Fig. S4B** and **Supplementary Video 1**). Furthermore, we were able to partially determine the specific location of PI4P on the surface area of the lysosome with Opto-PI4PSps (**Fig. S4B**). It has been shown that PI4P would be transferred from lysosomes to mitochondria at their interacting site^32^. Through the Opto-PI4PSps biosensor, we could readily observe that some intracellular PI4P was localized at the interaction site between mitochondria and lysosomes (**Fig. S4B** and **Supplementary Video 1**), suggesting its potential application for tracking phospholipids transfer at organelle interaction sites. These advantages were not afforded by traditional PI4P biosensors (**Fig. S4A**).

**Figure 4.**
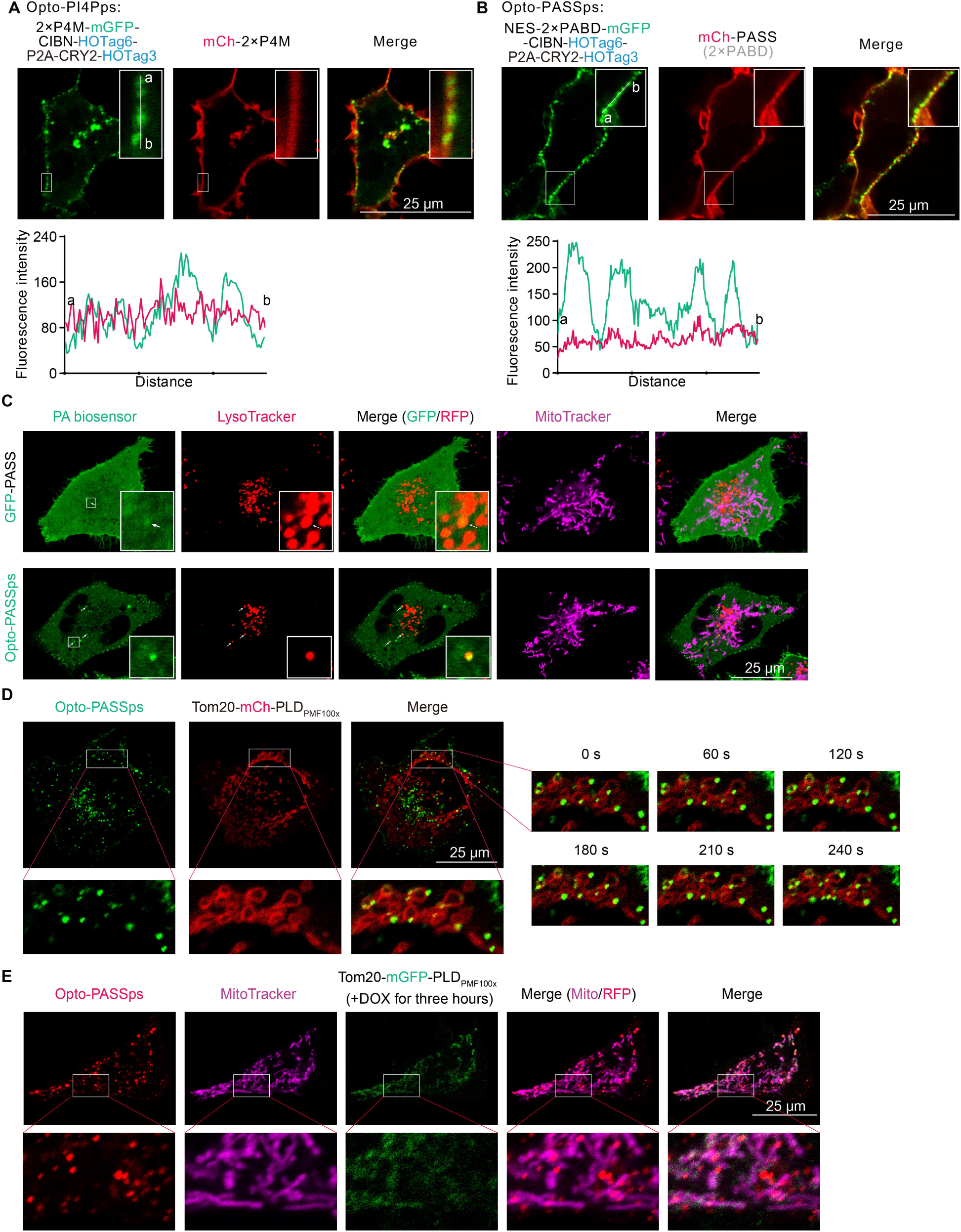
the performance of Opto-PI4PSps and Opto-PASSps in detecting intracellular phospholipids. **A**, a comparison of detecting performance of Opto-PI4PSps and Opto-PASSps (**B**) with the corresponding non-optimized biosensor after 10-minute blue light illumination, respectively; the fluorescence intensity of the white line (ab) was analyzed to assess the amplified signals. Using Opto-PASSps to detect basal levels of PA on the mitochondria and lysosomes (**C**) and visualize intracellular PA dynamics after overexpression of highly-active PLD_PMF100×_ on the mitochondria (**D**) in HeLa cells. **E**, detecting nascent PA production mediated by Dox-induced PLD_PMF100×_ expression with Opto-PASSps.

Due to the relatively low levels of PA under basal conditions and the interference from unbound biosensors, the non-optimized PASS sensor encountered challenges in detecting small amounts of PA on specific organelles. Consequently, we cautiously speculated about the presence of PA on lysosomes through co-localization analysis (**Fig. 4C**). However, this situation could be improved by using Opto-PASSps for detection. PA could be visually observed to co-localize with lysosomes and exhibit synchronous movement along with them through our improved biosensor (**Fig. 4C** and **Supplementary Video 2**). This observation was consistent with findings using fluorescent dye to label PA, such as IMPACT^25^. Furthermore, it has been shown that the proportion of PA in total phospholipids is less than 1% on the mitochondrial membrane^33^. In our results, co-localizations of punctum-like PA biosensors with mitochondria were scarcely observed (**Fig. 4C**), further suggesting a low basal level of PA on the mitochondrial membrane. Increasing PA has been implicated in mitochondrial dynamics and mitophagy^34-37^ and addressing its spatiotemporal regulatory mechanisms required high-performance biosensor. To elevate the PA level on the mitochondria, we overexpressed mitochondria-localized PLD_PMF100×_, resulting in substantial puncta located on the outer membrane of mitochondria as detected by both GPF-fused and mCh-fused Opto-PASSps biosensor (**Fig. 4D** and **Fig. S4C**). We also employed DOX to induce PLD_PMF100×_ expression and observed that our enhanced biosensor could detect the PA on the mitochondria induced by low levels of PLD expression (**Fig. 4E**), indicating its high affinity for PA and superior performance in detecting minor PA changes.

Currently, the SunTag system is a powerful tool for amplifying signals by fusion of multicopy peptides into the target molecule. These short peptides could be recognized by the single-chain variable fragments (scFV) antibody or nanoantibody. The resulting clusters resemble punctum-like shapes that amplify single-molecular signals^14^. In this system, it is crucial to control the number of fluorescent clusters to prevent a significant amount of dissociated probes from affecting the visualization of puncta. Therefore, we commonly used tetracycline to induce the expression of the target molecule. To investigate the potential of the SunTag system in enhancing biosensor performance, we fused two copies of PABDs with the tandem GCN4 peptides (24×GCN4) (here referred to as **PASS** based on the **S**un**T**ag system (PASSst)) and co-expressed them with anti-GCN4 scFV antibody probes in HeLa cells. After the addition of DOX, punctum-like structures were clearly observed (**Fig. 5A**). These puncta exhibited a size similar to the aforementioned optogenetic biosensors but displayed a lower brightness ratio (the ratio of spot brightness to the background) (**Fig. 5A**). We co-expressed PASSst and Opto-PASSps in cells overexpressing Tom20-PLD_PMF100_ ×. The red Opto-PASSps spots were partially colocalized with green punctate PASSst, and some of these puncta were located on the mitochondria (**Fig. 5B** □ cyan arrows), which suggests PLD-induced PA production and also indicates their spatial location on the mitochondria. However, a small set of the colocalized PA biosensors (yellow spots) were not on the mitochondria and moved faster than mitochodrial spots in cells (**Fig. 5B** ③ yellow arrows). As mentioned earlier, there were undeniable levels of PA on the lysosomes, leading us to suspect that these spots might label the PA present on the lysosomes.

**Figure 5.**
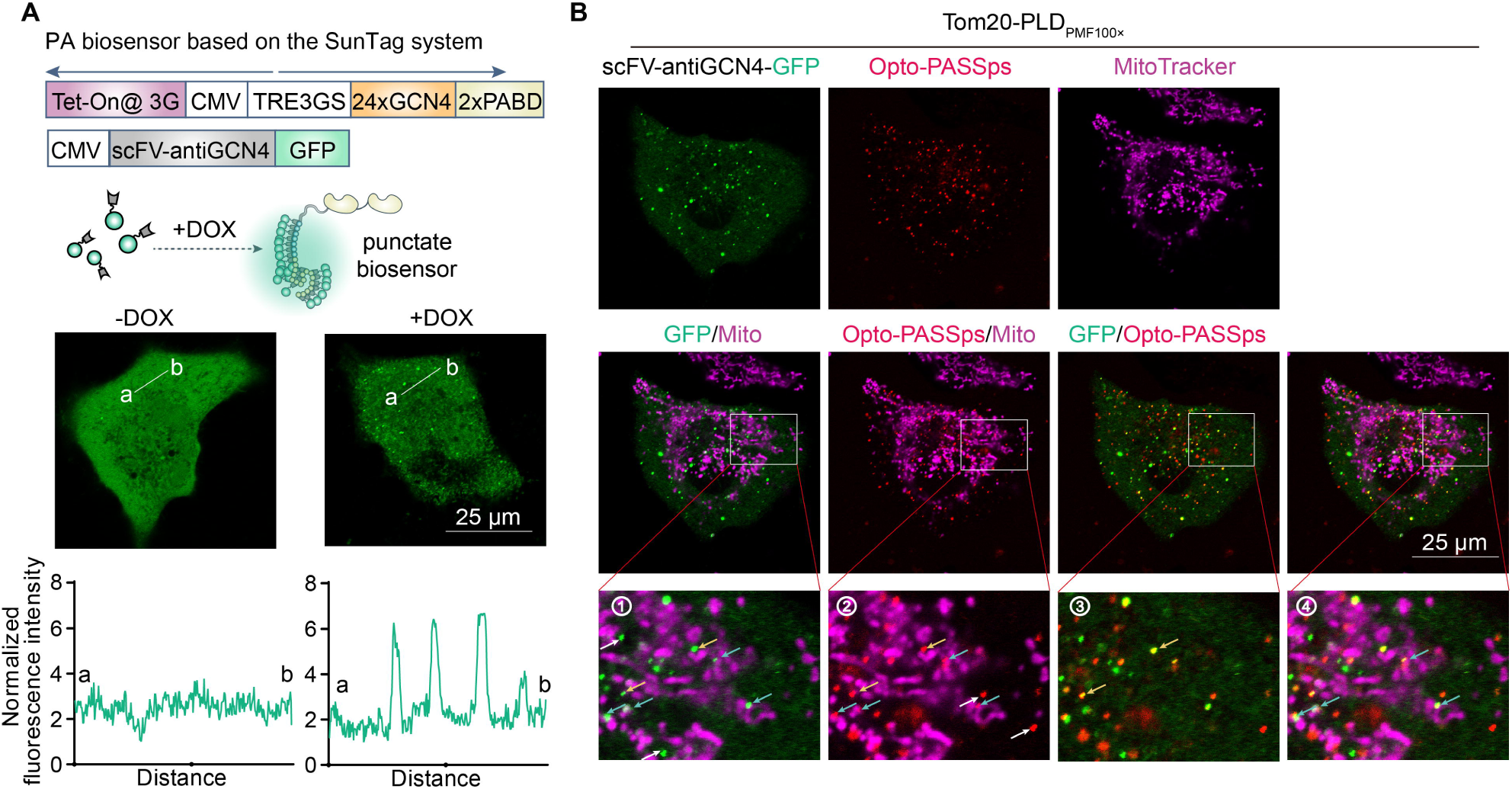
improving the PA biosensor based on the SunTag system and its comparison with Opto-PASSps. **A**, using the SunTag system to improve the PA biosensor, the signal of the line (ab) was normalized to the minimum fluorescence intensity in the line (ab); **B**, the PA-detecting performance of PASSst and Opto-PASSps in PLD-mediated PA production on the mitochondria, cyan arrow: co-localized PASSst and Opto-PASSps on the mitochondria, yellow arrow: co-localized PASSst and Opto-PASSps not on the mitochondria, white arrow in L: non-colocalized green PASSst biosensor, white arrow in L: non-colocalized red Opto-PASSps biosensor.

Taken together, the strategy of optically-controlled phase separation generated a similar amplification of the fluorescence signal of the biosensors in comparison to the SunTag system, but distinguished themselves by their light-controllable puncta formation rather than drug-induced expression. These punctum-like structures allowed us to visualize the intracellular distribution of PA and provided convenience to investigate the spatiotemporal regulatory roles of phospholipids in live cells. Additionally, the intense brightness of both punctate biosensors also provided protection against photobleaching during prolonged exposure, unlike traditional biosensors. The presence of multiple copies of phospholipid-binding motifs within each punctum, induced by phase separation, conferred greater sensitivity to these engineering biosensors.

## 3 Conclusion

In summary, our study represents a significant advancement in utilizing optogenetic strategies to enhance the functionality of phospholipid biosensors. Compared to traditional approaches in biosensor development, these optogenetics-based enhancements effectively reduce background signals, providing clearer reflections of phospholipid changes in cells. The application of optically-controlled phase separation induces the formation of fluorescent spots, amplifying signals and facilitating phospholipids visualization in cells. The improved biosensors presented in our study enriched the toolbox for phospholipid imaging and they hold great potential for enhancing our understanding of the spatio-temporal dynamics and regulatory roles of membrane lipids in living cells. The methodological improvements outlined in this study offer valuable insights for the development of other high-performance biosensors.

## 4 Experimental Section

### The construction of optogenetics-based biosensors

For CRY2-based sensors, the mcherry-CRY2 was amplified from the mCherry-mCh-CRY2-PR variant^38^. The sequence of mGFP-NES-PABP was amplified from GFP-PASS, a gift from Prof. Guangwei Du. These fragments were subcloned into pLenti-MCS-GFP-SV40-Puro empty vector that linearized by XbaI and SpeI restriction enzymes (New England Biolabs, Inc., USA) via Gibson Assembly kit (TransGen Biotech., China).

For cpLOV2-based sensor, we first synthesized cpLOV2 sequence of which the Jα helix contained a NES sequence. To replace different NES, we inserted two BsmBI sites into both ends of NES. cpLOV2, together with mCherry-NLS, was then cloned into linearized pLenti-MCS-GFP-SV40-Puro empty vector. This framework (pLenti-cpLOV2-mCherry-NLS) was further modified by the replacement of mCherry-NLS with NLS-mGFP-NES-PABP, namely Opto-PASSc in this study. We also constructed red Opto-PASSc probe by the replacement of mGFP with mCherry sequence. PABP 4E mutations were introduced into the PABP sequence by primers and its PCR fragment, together with cpLOV2-NLS-mCherry-NES fragment, was inserted into corresponding linearized pLenti-MCS-GFP-SV40-Puro.

The PI4P sensor based on optically-controlled phase separation was created by fusion of P4M domian from mCherry-P4M-SidM (#51471, Addgene, USA) to mGFP-CIBN. The mGFP (Emerald) was amplified from Tom20-Emerald (#54282, Addgene, USA) and the CIBN was amplified from CIBN-CAAX (#79574, Addgene, USA). The mCherry-CRY2 was amplified as described previously. HOTag3 and HOTag6 was synthesized (Beijing Tsingke Biotech Co., Ltd. China) with corresponding homologous sequence in both ends and a sequence encoding a specific linker. They were subcloned into linearized vector with (P4M)_n_-mGFP-CIBN and mCh-CRY2, respectively. The PLD_PMF100×_ was synthesized (Beijing Tsingke Biotech Co., Ltd. China) according to Baskin’s publication^30^. The PASSst sensor was created by fusion of two copies of PABD with the GCN4 that was amplified from pcDNA4TO-24xGCN4_v4-kif18b (#74934, Addgene, USA). The GCN4 probe was pHR-scFv-GCN4-sfGFP-GB1-dWPRE (#60907, Addgene, USA). The above plasmids were assembled with the corresponding purified PCR fragments via Gibson Assembly kit. The sequence and source of other constructions used in this study are listed in Supplementary Materials and all plasmids used in this study were sequenced for validation.

### The cell culture and transfection

Cancer cell lines HeLa and 293T used in our research were purchased from ATCC. 293T cell were routinely grown in Dulbecco’s Modified Essential Media (DMEM, Hyclone), while HeLa cells were routinely grown in RPMI 1640. The complete medium was supplemented with 10% fetal bovine serum (PAN, PAN-Biotech, Germany) and 100 U/ml penicillin/streptomycin (Beyotime Biotechnology, China). The cells were cultured at 37 °C in a humidified atmosphere with 5% CO_2_. For transient over-expression assay, the cells were plated on confocal dishes. About 1-2 μg indicated plasmid was used according to the instruction of lipofectamine 3000 (Thermo Fisher Scientific, USA).

### Living cell imaging

Cells were plated on the confocal dish and then were co-transfected with indicated plasmid. After 24-hour transfection, refreshed with the phenol-free medium. The confocal images were obtained using a AX confocal laser scanning microscope (Nikon, Japan) equipped with a 60×/1.49 numerical aperture (NA) oil immersion objective lens. For light-induced PA probe export, the blue light pulse was 10-12 seconds and the 488 laser power was 1 mW. For the time-scale detection of PA at plasma membrane, such as PMA-induced PA production, the first image was captured under non-stimulation condition and subsequently 10 μL PBS buffer containing 500 μM PMA (P8139, Sigma-Aldrich, USA) was added into the dish carefully. The capture interval was 8 minutes. For time-scale detection of PI4P at plasma membrane, the images were captured by a C2 confocal laser scanning microscope with a TIRF module (Nikon, Japan). The first image was captured and subsequently PAO (P3075, Sigma-Aldrich, USA) with the indicated final concentration was added into the dish carefully. The capture interval was 10 minutes.

### Statistical analysis

The fluorescence intensity of the images was analyzed by FiJi Image J software (Version 1.53t). Two experimental groups were analyzed by Student’s *t*-test (one-tail test). Multigroup comparisons were performed by the one-way ANOVA with Tukey’s post hoc test. The *p* values less than 0.05 were considered as significant. Replicates consist of at least three independent experiments. Quantitative data are presented as mean ± SEM. GraphPad Prism were used for data analysis and plot.

## Acknowledgements

We thank Prof. Guangwei Du for the gifts RFP-PASS and GFP-PASS. This research was supported by the National key research and development program (2021YFC2101001), the Cross Scientific Research Fund of Zhejiang University of Technology (2022JCY27) and the Natural Science Foundation of Zhejiang Province (LY18H070004).

## Author Contributions

Y.Y, X.L., and L.H. conceived the idea of optogenetic strategies for improvement of biosensors and discussed the optimization of opto-biosensors. H.L., L.H. and Y.X. supervised the research. Y.Y, and X.L. designed and conducted most of the experiments. J.L., W.S., Y.C., S.C, T.Z., and S.K. prepared the samples. Y.Y, X.L., J.L., W.S., and Y.C. analyzed the experimental data and plotted figures. Y.Y, H.L., L.H., and Y.X. wrote and revised the paper with input from all authors.

## References

1. He, L., Tan, P., Zhu, L., Huang, K., Nguyen, N.T., Wang, R., Guo, L., Li, L., Yang, Y., and Huang, Z. (2021). Circularly permuted LOV2 as a modular photoswitch for optogenetic engineering. Nature Chemical Biology 17, 915–923.

2. Yumerefendi, H., Lerner, A.M., Zimmerman, S.P., Hahn, K., Bear, J.E., Strahl, B.D., and Kuhlman, B. (2016). Light-induced nuclear export reveals rapid dynamics of epigenetic modifications. Nature Chemical Biology 12, 399–401. 10.1038/nchembio.2068.

3. Strickland, D., Lin, Y., Wagner, E., Hope, C.M., Zayner, J., Antoniou, C., Sosnick, T.R., Weiss, E.L., and Glotzer, M. (2012). TULIPs: tunable, light-controlled interacting protein tags for cell biology. Nature methods 9, 379–384.

4. Guntas, G., Hallett, R.A., Zimmerman, S.P., Williams, T., Yumerefendi, H., Bear, J.E., and Kuhlman, B. (2015). Engineering an improved light-induced dimer (iLID) for controlling the localization and activity of signaling proteins. Proceedings of the National Academy of Sciences 112, 112–117.

5. Liu, R., Yang, J., Yao, J., Zhao, Z., He, W., Su, N., Zhang, Z., Zhang, C., Zhang, Z., Cai, H., et al. (2022). Optogenetic control of RNA function and metabolism using engineered light-switchable RNA-binding proteins. Nature Biotechnology 40, 779–786. 10.1038/s41587-021-01112-1.

6. Kennedy, M.J., Hughes, R.M., Peteya, L.A., Schwartz, J.W., Ehlers, M.D., and Tucker, C.L. (2010). Rapid blue-light–mediated induction of protein interactions in living cells. Nature Methods 7, 973–975. 10.1038/nmeth.1524.

7. He, L., Wang, L., Zeng, H., Tan, P., Ma, G., Zheng, S., Li, Y., Sun, L., Dou, F., and Siwko, S. (2021). Engineering of a bona fide light-operated calcium channel. Nature Communications 12, 164.

8. Li, T., Chen, X., Qian, Y., Shao, J., Li, X., Liu, S., Zhu, L., Zhao, Y., Ye, H., and Yang, Y. (2021). A synthetic BRET-based optogenetic device for pulsatile transgene expression enabling glucose homeostasis in mice. Nature Communications 12, 615.

9. Zhou, Y., Kong, D., Wang, X., Yu, G., Wu, X., Guan, N., Weber, W., and Ye, H. (2022). A small and highly sensitive red/far-red optogenetic switch for applications in mammals. Nature Biotechnology 40, 262–272. 10.1038/s41587-021-01036-w.

10. Yu, Y., Wu, X., Guan, N., Shao, J., Li, H., Chen, Y., Ping, Y., Li, D., and Ye, H. (2020). Engineering a far-red light–activated split-Cas9 system for remote-controlled genome editing of internal organs and tumors. Science Advances 6, eabb1777. doi:10.1126/sciadv.abb1777.

11. Deisseroth, K., and Hegemann, P. (2017). The form and function of channelrhodopsin. Science 357, eaan5544. doi:10.1126/science.aan5544.

12. Duan, L., Che, D., Zhang, K., Ong, Q., Guo, S., and Cui, B. (2015). Optogenetic Control of Molecular Motors and Organelle Distributions in Cells. Chemistry & Biology 22, 671–682. 10.1016/j.chembiol.2015.04.014.

13. van Bergeijk, P., Adrian, M., Hoogenraad, C.C., and Kapitein, L.C. (2015). Optogenetic control of organelle transport and positioning. Nature 518, 111–114. 10.1038/nature14128.

14. Tanenbaum, M.E., Gilbert, L.A., Qi, L.S., Weissman, J.S., and Vale, R.D. (2014). A protein-tagging system for signal amplification in gene expression and fluorescence imaging. Cell 159, 635–646. 10.1016/j.cell.2014.09.039.

15. Wang, C., Han, B., Zhou, R., and Zhuang, X. (2016). Real-Time Imaging of Translation on Single mRNA Transcripts in Live Cells. Cell 165, 990–1001. 10.1016/j.cell.2016.04.040.

16. Yan, X., Hoek, Tim A., Vale, Ronald D., and Tanenbaum, Marvin E. (2016). Dynamics of Translation of Single mRNA Molecules In Vivo. Cell 165, 976–989. 10.1016/j.cell.2016.04.034.

17. Zhao, N., Kamijo, K., Fox, P.D., Oda, H., Morisaki, T., Sato, Y., Kimura, H., and Stasevich, T.J. (2019). A genetically encoded probe for imaging nascent and mature HA-tagged proteins in vivo. Nature Communications 10. 10.1038/s41467-019-10846-1.

18. Boersma, S., Khuperkar, D., Verhagen, B.M.P., Sonneveld, S., Grimm, J.B., Lavis, L.D., and Tanenbaum, M.E. (2019). Multi-Color Single-Molecule Imaging Uncovers Extensive Heterogeneity in mRNA Decoding. Cell 178, 458–472.e419. 10.1016/j.cell.2019.05.001.

19. Zhang, Q., Huang, H., Zhang, L., Wu, R., Chung, C.I., Zhang, S.Q., Torra, J., Schepis, A., Coughlin, S.R., Kornberg, T.B., and Shu, X. (2018). Visualizing Dynamics of Cell Signaling In Vivo with a Phase Separation-Based Kinase Reporter. Mol Cell 69, 334–346.e334. 10.1016/j.molcel.2017.12.008.

20. Li, X., Combs, J.D., Salaita, K., and Shu, X. (2023). Polarized focal adhesion kinase activity within a focal adhesion during cell migration. Nature Chemical Biology 19, 1458–1468. 10.1038/s41589-023-01353-y.

21. Li, X., Chung, C.-I., Yang, J., Chaudhuri, S., Munster, P.N., and Shu, X. (2023). ATM-SPARK: A GFP phase separation–based activity reporter of ATM. Science Advances 9, eade3760. doi:10.1126/sciadv.ade3760.

22. Yao, Y., Li, H., Wang, X., Sun, Y., Zhao, X., Zha, W., Zhou, J., Toomre, D., Fu, J., and Xu, Y. (2023). Novel function of biguanides in inhibition of phospholipase D1 expression via a translational mechanism in cancer cells. Genes Dis 10, 1787–1790. 10.1016/j.gendis.2023.01.007.

23. Yao, Y., Li, J., Lin, Y., Zhou, J., Zhang, P., and Xu, Y. (2021). Structural insights into phospholipase D function. Progress in Lipid Research 81, 101070. 10.1016/j.plipres.2020.101070.

24. Yao, Y., Wang, X., Li, H., Fan, J., Qian, X., Li, H., and Xu, Y. (2020). Phospholipase D as a key modulator of cancer progression. Biological Reviews 95, 911–935. 10.1111/brv.12592.

25. Bumpus, T.W., and Baskin, J.M. (2017). Clickable substrate mimics enable imaging of phospholipase D activity. ACS central science 3, 1070–1077.

26. Liang, D., Wu, K., Tei, R., Bumpus, T.W., Ye, J., and Baskin, J.M. (2019). A real-time, click chemistry imaging approach reveals stimulus-specific subcellular locations of phospholipase D activity. Proc Natl Acad Sci U S A 116, 15453–15462. 10.1073/pnas.1903949116.

27. Bumpus, T.W., Huang, S., Tei, R., and Baskin, J.M. (2021). Click chemistry-enabled CRISPR screening reveals GSK3 as a regulator of PLD signaling. Proc Natl Acad Sci U S A 118. 10.1073/pnas.2025265118.

28. Zhang, F., Wang, Z., Lu, M., Yonekubo, Y., Liang, X., Zhang, Y., Wu, P., Zhou, Y., Grinstein, S., Hancock, J.F., and Du, G. (2014). Temporal Production of the Signaling Lipid Phosphatidic Acid by Phospholipase D2 Determines the Output of Extracellular Signal-Regulated Kinase Signaling in Cancer Cells. Molecular and Cellular Biology 34, 84–95. 10.1128/MCB.00987-13.

29. Yao, Y., Li, H., Wang, X., Sun, Y., Zhao, X., Zha, W., Zhou, J., Toomre, D., Fu, J., and Xu, Y. (2023). Novel function of biguanides in inhibition of phospholipase D1 expression via a translational mechanism in cancer cells. Genes & Diseases 10, 1787–1790. 10.1016/j.gendis.2023.01.007.

30. Tei, R., Bagde, S.R., Fromme, J.C., and Baskin, J.M. (2023). Activity-based directed evolution of a membrane editor in mammalian cells. Nature Chemistry 15, 1030-1039. 10.1038/s41557-023-01214-0.

31. Taslimi, A., Zoltowski, B., Miranda, J.G., Pathak, G.P., Hughes, R.M., and Tucker, C.L. (2016). Optimized second-generation CRY2–CIB dimerizers and photoactivatable Cre recombinase. Nature Chemical Biology 12, 425–430. 10.1038/nchembio.2063.

32. Boutry, M., and Kim, P.K. (2021). ORP1L mediated PI(4)P signaling at ER-lysosome-mitochondrion three-way contact contributes to mitochondrial division. Nat Commun 12, 5354. 10.1038/s41467-021-25621-4.

33. Horvath, S.E., and Daum, G. (2013). Lipids of mitochondria. Prog Lipid Res 52, 590–614. 10.1016/j.plipres.2013.07.002.

34. Lin, C.C., Yan, J., Kapur, M.D., Norris, K.L., Hsieh, C.W., Huang, D., Vitale, N., Lim, K.L., Guan, Z., Wang, X.F., et al. (2022). Parkin coordinates mitochondrial lipid remodeling to execute mitophagy. EMBO reports 23, e55191. 10.15252/embr.202255191.

35. Nagdas, S., Kashatus, J.A., Nascimento, A., Hussain, S.S., Trainor, R.E., Pollock, S.R., Adair, S.J., Michaels, A.D., Sesaki, H., Stelow, E.B., et al. (2019). Drp1 Promotes KRas-Driven Metabolic Changes to Drive Pancreatic Tumor Growth. Cell Reports 28, 1845–1859.e1845. 10.1016/j.celrep.2019.07.031.

36. Choi, S.-Y., Huang, P., Jenkins, G.M., Chan, D.C., Schiller, J., and Frohman, M.A. (2006). A common lipid links Mfn-mediated mitochondrial fusion and SNARE-regulated exocytosis. Nature Cell Biology 8, 1255–1262. 10.1038/ncb1487.

37. Perea, V., Cole, C., Lebeau, J., Dolina, V., Baron, K.R., Madhavan, A., Kelly, J.W., Grotjahn, D.A., and Wiseman, R.L. (2023). PERK signaling promotes mitochondrial elongation by remodeling membrane phosphatidic acid. The EMBO Journal 42. 10.15252/embj.2023113908.

38. Tan, P., Hong, T., Cai, X., Li, W., Huang, Y., He, L., and Zhou, Y. (2022). Optical control of protein delivery and partitioning in the nucleolus. Nucleic Acids Research 50, e69–e69.

