## Supplementary Materials for "Optogenetic strategies for optimizing the performance of biosensors of membrane phospholipids in live cells"

1. **Supplementary figures (1-4)**

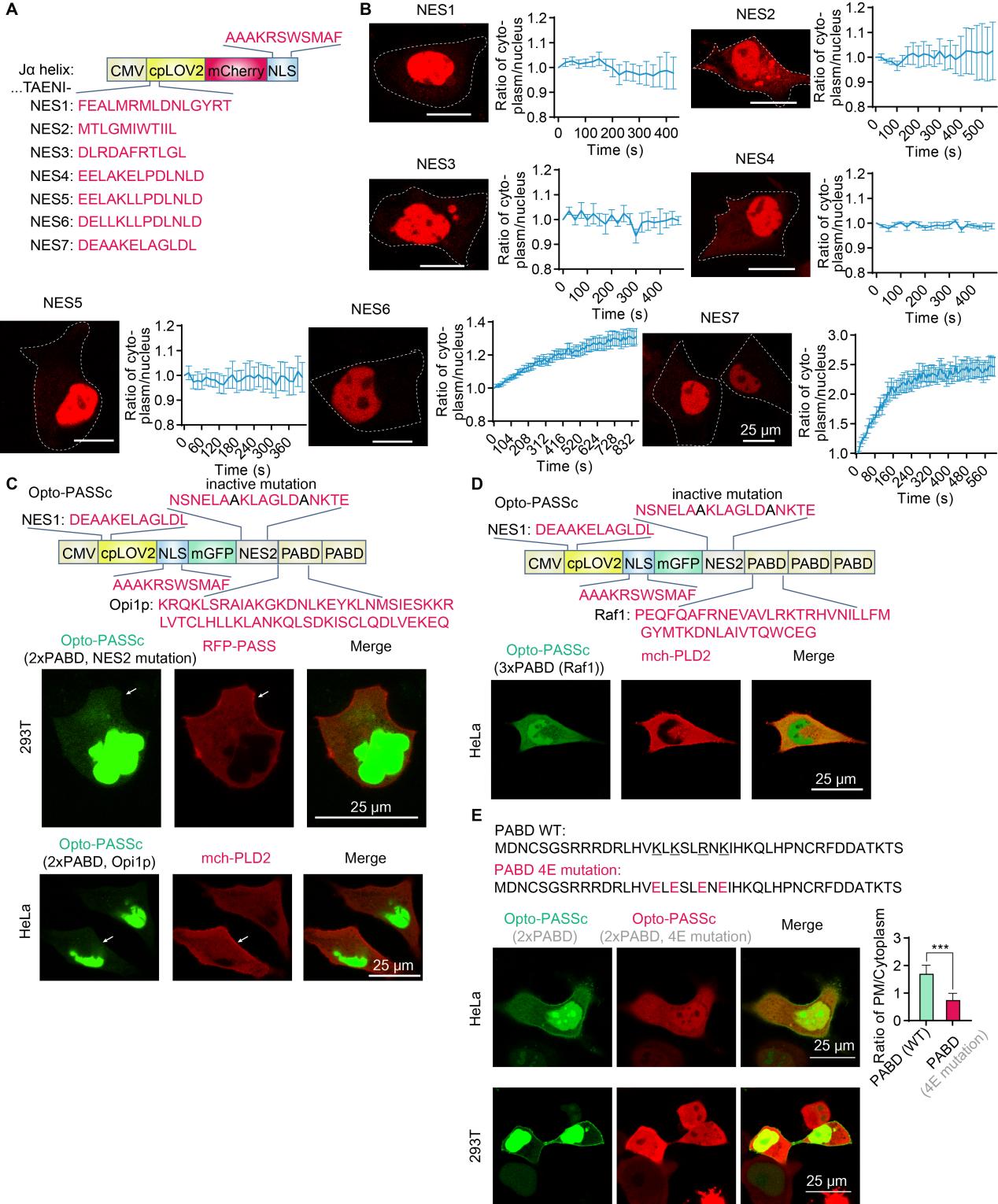
**Supplementary figure 1** **screening of NES for regulating the nuclear export of cpLOV2 and evaluation of the PA binding motif among various PA binding proteins. A**, the design of cpLOV-based nucleocytoplasmic transportation and **B**, representative images of cpLOV-mCh-NLS distribution in cells under non-stimulation condition and quantitative analysis of the light-induced nuclear export of cpLOV-mCh-NLS with fusion of different NES sequences. **C**, the schematic diagram of the cpLOV-based PA biosensor (Opto-PASSc) with two PABD motifs from yeast Opi1p and representative images of co-expression of Opto-PASSc with RFP-PASS or mCh-PLD2 in HeLa cells and in 293T cells. **D**, the schematic diagram of the cpLOV-based PA biosensor (Opto-PASSc) with three PABD motifs from Raf1 protein and representative images of its co-expression with mCh-PLD2. **E**, amino acid sequence of PABD and 4E mutant PABD and representative images of Opto-PASSc with or without 4E mutation in HeLa cells and in 293T cells. The ratio of plasma membrane to cytosol were analyzed in HeLa (n=11 cells).

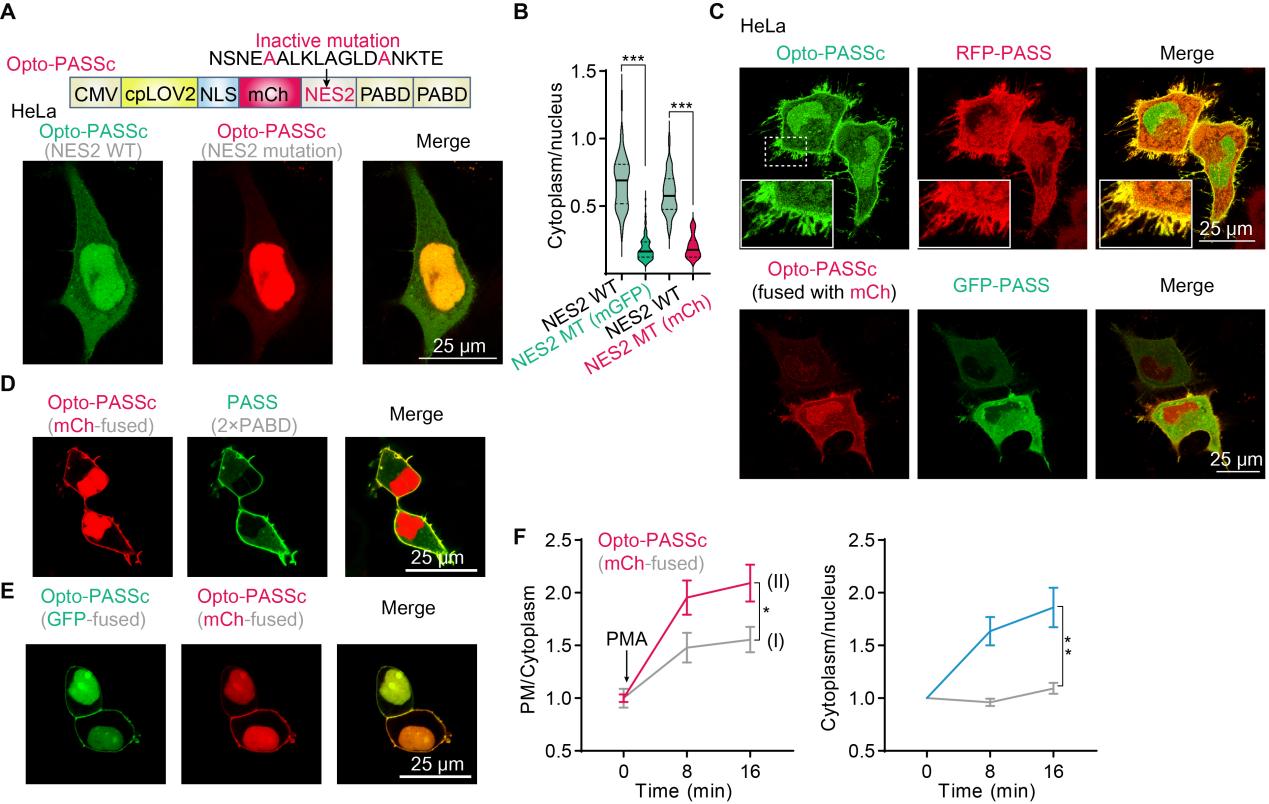

**Supplementary figure 2 Optimization of Opto-PASSc in** **background noise and its comparison with the wildly-used PASS. A**, further reducing the background noise from the unbound PASS by inactive mutation of internal NES2, two residues (Leucine and Isoleucine) of the NES2 was mutated into Alanine and colored in red. **B**, assessing the improvement of the background signal after NES2 mutations by measuring the fluorescence intensity ratio of the cytosol to the nucleus , this analysis involved comparing the following groups: wild-type NES2 vs. NES2 with mutations (mGFP-fused) (n ≥ 70 cells) and wild-type NES2 vs. NES2 with mutations (mCherry-fused) (n = 27 cells). **C**, the colocalization of Opto-PASSc (fused with mGFP or mCherry) with the wildly-used PASS biosensor in HeLa cells. **D**, representative images of co-expression of the mCh-fused Opto-PASSc with the GFP-PASS containing the same copy of PABD, and **E**, representative images of co-expression of Opto-PASSc (2×PABD) fused with GFP or mCh. **F**, the impact of blue light stimulation on the PA-detecting performance of mCh-fused Opto-PASSc in response to PMA (5 μM)-induced PA production, the condition (I): only three images were captured at indicated time and the interval was dark treatment; the condition (II): blue light stimulation at the capture interval, PMA was added after finishing capture of the first image, the data was normalized to the initial value (n = 7 cells). The PM/Cytoplasm ratio was normalized to the mean of the initial value, and the data of Cytoplasm/nucleus was normalized to their respective initial values.

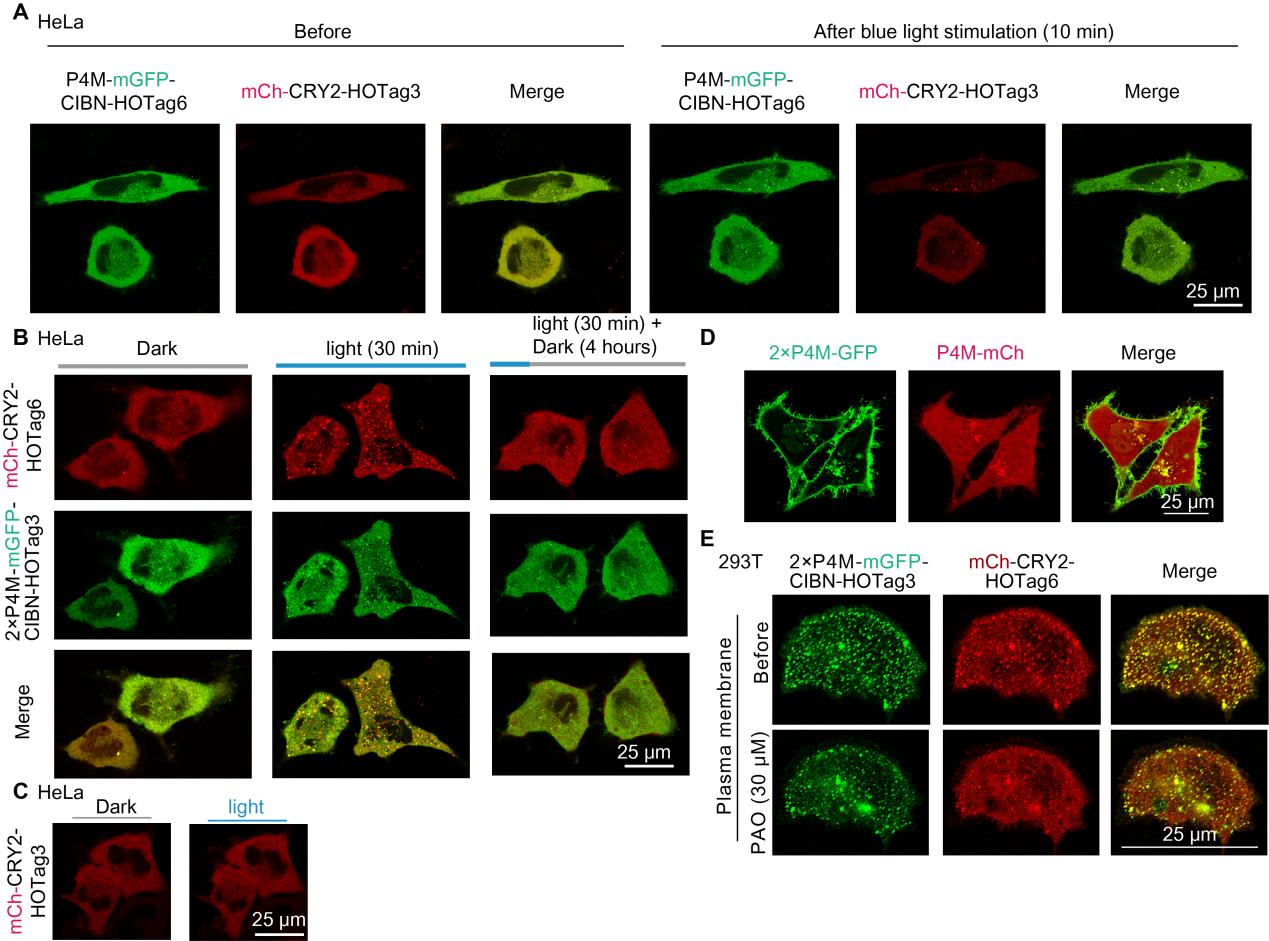
**Supplementary figure 3 the optically-controlled phase separation amplified the signal of PI4P biosensor and its application in detecting PI4P changes after chemical inhibitor treatment.** **A**, the light-induced binding of CRY2/CIBN triggered the phase separation via HOTag3/HOTag6 in HeLa cells with co-expression of indicated plasmids. **B**, the disaggregation of the light-induced phase separation after dark treatment. **C**, the phase separation did not induced by light in the cells with single expression of mCh-CRY2-HOTag3. **D**, the representative image of PI4P biosensors with one or two copies of P4M. **E**, the enhanced probe Opto-PI4PSps was employed to monitor PI4P changes on the plasma membrane in the cells in respond to a half-hour treatment with a PI4P inhibitor, PAO (30 μM).

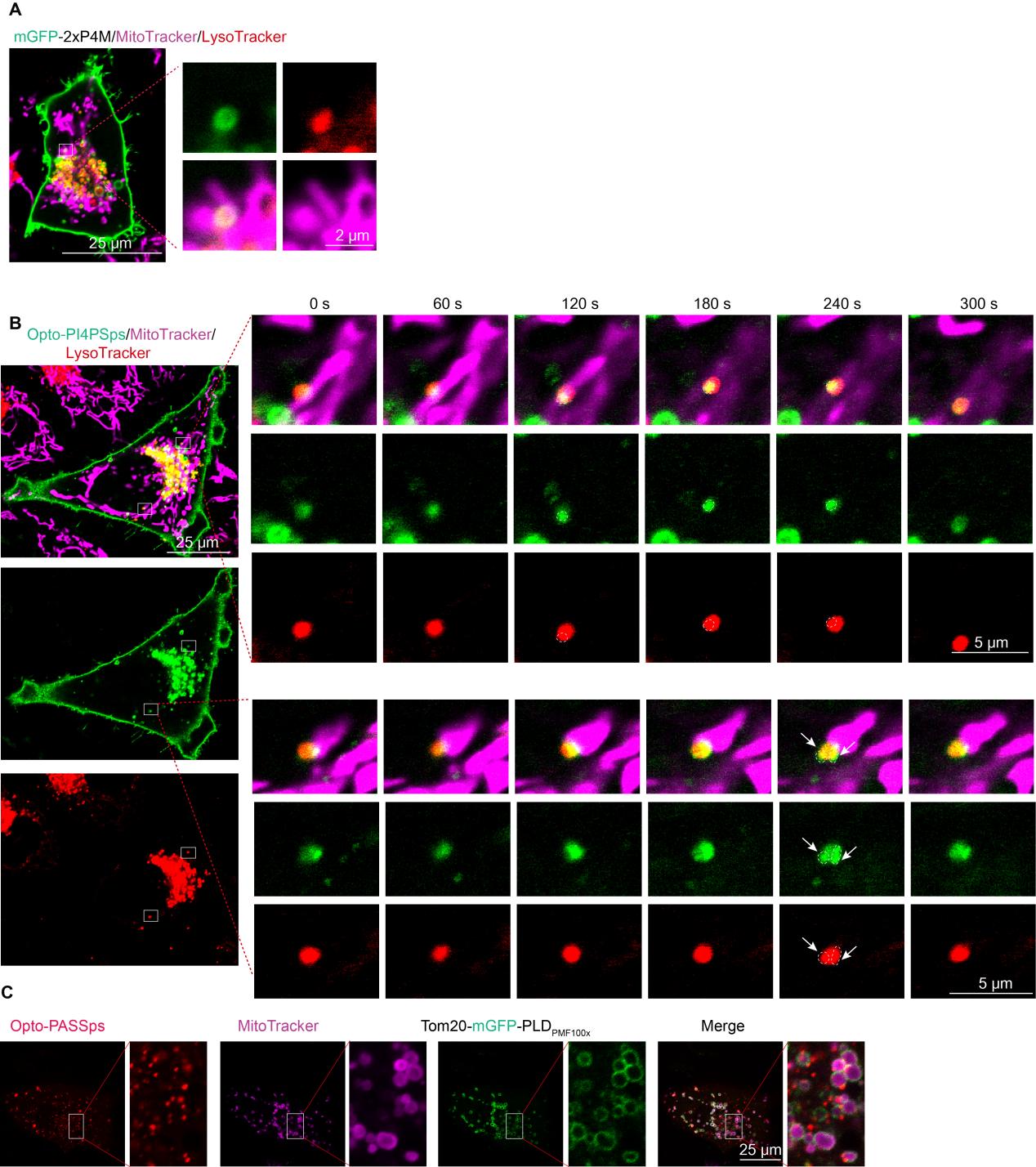

**Supplementary figure 4** a comparison of mGFP-2×P4M (**A**) with Opto-PI4PSps (**B**) in visualizing intracellular PI4P. PI4P was localized on some of lysosomes through the non-optimized PI4P biosensor, while Opto-PI4PSps visually and informatively presented PI4P on the lysosomes and the interacting site of lysosome and mitochondria. **C**, Using mCh-fused Opto-PASSps to visualize intracellular PA dynamics after overexpression of highly-active PLD_PMF100×_ on the mitochondria.

**2. sequence of plasmids used in this study.**

| Plasmid | Sequence |
| --- | --- |
| pLenti-CMV-cpLOV2(NES1 in the Jα helix)  -NLS-mGFP(or mCh)  -NES2(MT or WT)-  2×PABD(or 4×PABD) | cpLOV2: yellow linker: red NLS: blue   mGFP: green  NES1: gray PABP: deep yellow NES2: gray  MTEHVRDAAEREGVMLIKKTAENIDEAAKELAGLDLGGGSGGSGGGLATTLERIEKNFVITDPRLPDNPIIFASDSFLQLTEYSREEILGRNCRFLQGPETDRATVRKIRDAIDNQTEVTVQLINYTKSGKKFWNLFHLQPMRDQKGDVQYFIGVQLDGGSGSGSAAAKRSWSMAFGSGSGSMVSKGEELFTGVVPILVELDGDVNGHKFSVSGEGEGDATYGKLTLKFICTTGKLPVPWPTLVTTLTYGVQCFSRYPDHMKQHDFFKSAMPEGYVQERTIFFKDDGNYKTRAEVKFEGDTLVNRIELKGIDFKEDGNILGHKLEYNYNSHNVYIMADKQKNGIKVNFKIRHNIEDGSVQLADHYQQNTPIGDGPVLLPDNHYLSTQSKLSKDPNEKRDHMVLLEFVTAAGITLGMDELYKSGLRSRANSNE**A**ALKLAGLD**A**NKTESRMDNCSASRRRDRLHVKLKSLRNKIHKQLHPNCRFDDATKTSGGGSGGGSMDNCSGSRRRDRLHVKLKSLRNKIHKQLHPNCRFDDATKTS  **NES2WT:** NSNELALKLAGLDINKTE  **mCherry:** MVSKGEEDNMAIIKEFMRFKVHMEGSVNGHEFEIEGEGEGRPYEGTQTAKLKVTKGGPLPFAWDILSPQFMYGSKAYVKHPADIPDYLKLSFPEGFKWERVMNFEDGGVVTVTQDSSLQDGEFIYKVKLRGTNFPSDGPVMQKKTMGWEASSERMYPEDGALKGEIKQRLKLKDGGHYDAEVKTTYKAKKPVQLPGAYNVNIKLDITSHNEDYTIVEQYERAEGRHSTGGMDELYK  **4×PABD:**  MDNCSASRRRDRLHVKLKSLRNKIHKQLHPNCRFDDATKTSGGGSGGGSMDNCSGSRRRDRLHVKLKSLRNKIHKQLHPNCRFDDATKTSGSGSGSGSANKTESRMDNCSASRRRDRLHVKLKSLRNKIHKQLHPNCRFDDATKTSGGGSGGGSMDNCSGSRRRDRLHVKLKSLRNKIHKQLHPNCRFDDATKTS |
| pLenti-mGFP(mCh)-NES2(WT)-2×PABD (or 4×PABD) | linker: red NLS: blue mGFP: green NES2: deep gray PABP: deep yellow  MVSKGEELFTGVVPILVELDGDVNGHKFSVSGEGEGDATYGKLTLKFICTTGKLPVPWPTLVTTLTYGVQCFSRYPDHMKQHDFFKSAMPEGYVQERTIFFKDDGNYKTRAEVKFEGDTLVNRIELKGIDFKEDGNILGHKLEYNYNSHNVYIMADKQKNGIKVNFKIRHNIEDGSVQLADHYQQNTPIGDGPVLLPDNHYLSTQSKLSKDPNEKRDHMVLLEFVTAAGITLGMDELYKSGLRSRANSNELALKLAGLDINKTESRMDNCSASRRRDRLHVKLKSLRNKIHKQLHPNCRFDDATKTSGGGSGGGSMDNCSGSRRRDRLHVKLKSLRNKIHKQLHPNCRFDDATKTS  **mCherry:** MVSKGEEDNMAIIKEFMRFKVHMEGSVNGHEFEIEGEGEGRPYEGTQTAKLKVTKGGPLPFAWDILSPQFMYGSKAYVKHPADIPDYLKLSFPEGFKWERVMNFEDGGVVTVTQDSSLQDGEFIYKVKLRGTNFPSDGPVMQKKTMGWEASSERMYPEDGALKGEIKQRLKLKDGGHYDAEVKTTYKAKKPVQLPGAYNVNIKLDITSHNEDYTIVEQYERAEGRHSTGGMDELYK  **4×PABD:**  MDNCSASRRRDRLHVKLKSLRNKIHKQLHPNCRFDDATKTSGGGSGGGSMDNCSGSRRRDRLHVKLKSLRNKIHKQLHPNCRFDDATKTSGSGSGSGSANKTESRMDNCSASRRRDRLHVKLKSLRNKIHKQLHPNCRFDDATKTSGGGSGGGSMDNCSGSRRRDRLHVKLKSLRNKIHKQLHPNCRFDDATKTS |
| pLenti-2×P4M-GFP-  CIBN-HOTag3 | P4M: deep yellow GFP: green linker: red CIBN: black linker-HOTag3: blue  MTASTENFKNVKEKYQQMRGDALKTEILADFKDKLAEATDEQSLKQIVAELKSKDEYRILAKGQGLTTQLLGLKTSSVSSFEKMVEETRESIKSQERQTIKIKGGGSGGGSTASTENFKNVKEKYQQMRGDALKTEILADFKDKLAEATDEQSLKQIVAELKSKDEYRILAKGQGLTTQLLGLKTSSVSSFEKMVEETRESIKSQERQTIKIKGGMVSKGEELFTGVVPILVELDGDVNGHKFSVSGEGEGDATYGKLTLKFICTTGKLPVPWPTLVTTLTYGVQCFARYPDHMKQHDFFKSAMPEGYVQERTIFFKDDGNYKTRAEVKFEGDTLVNRIELKGIDFKEDGNILGHKLEYNYNSHKVYITADKQKNGIKVNFKTRHNIEDGSVQLADHYQQNTPIGDGPVLLPDNHYLSTQSKLSKDPNEKRDHMVLLEFVTAAGITLGMDELYASGSGSMNGAIGGDLLLNFPDMSVLERQRAHLKYLNPTFDSPLAGFFADSSMITGGEMDSYLSTAGLNLPMMYGETTVEGDSRLSISPETTLGTGNFKAAKFDTETKDCNEAAKKMTMNRDDLVEEGEEEKSKITEQNNGSTKSIKKMKHKAKKEENNFSNDSSKVTKELEKTDYIHVGSGSAGGSAGGSAGGSAGGSAGGSAGGSAGGSRGEIAKSLKEIAKSLKEIAWSLKEIAKSLKG |
| pLenti-mCh-CRY2-  HOTag6 | mCherry: deep red CRY2: deep blue linker: red linker-HOTag6: blue  MVSKGEEDNMAIIKEFMRFKVHMEGSVNGHEFEIEGEGEGRPYEGTQTAKLKVTKGGPLPFAWDILSPQFMYGSKAYVKHPADIPDYLKLSFPEGFKWERVMNFEDGGVVTVTQDSSLQDGEFIYKVKLRGTNFPSDGPVMQKKTMGWEASSERMYPEDGALKGEIKQRLKLKDGGHYDAEVKTTYKAKKPVQLPGAYNVNIKLDITSHNEDYTIVEQYERAEGRHSTGGMDELYKRSRSAAAGAGGAARAMKMDKKTIVWFRRDLRIEDNPALAAAAHEGSVFPVFIWCPEEEGQFYPGRASRWWMKQSLAHLSQSLKALGSDLTLIKTHNTISAILDCIRVTGATKVVFNHLYDPVSLVRDHTVKEKLVERGISVQSYNGDLLYEPWEIYCEKGKPFTSFNSYWKKCLDMSIESVMLPPPWRLMPITAAAEAIWACSIEELGLENEAEKPSNALLTRAWSPGWSNADKLLNEFIEKQLIDYAKNSKKVVGNSTSLLSPYLHFGEISVRHVFQCARMKQIIWARDKNSEGEESADLFLRGIGLREYSRYICFNFPFTHEQSLLSHLRFFPWDADVDKFKAWRQGRTGYPLVDAGMRELWATGWMHNRIRVIVSSFAVKFLLLPWKWGMKYFWDTLLDADLECDILGWQYISGSIPDGHELDRLDNPALQGAKYDPEGEYIRQWLPELARLPTEWIHHPWDAPLTVLKASGVELGTNYAKPIVDIDTARELLAKAISRTREAQIMIGAAARGAAAGAGGAGRGGGGSGSGSAGGSAGGSAGGSAGGSAGGSAGGSAGGSRTLREIEELLRKIIEDSVRSVAELEDIEKWLKKI |
| pLenti-P4M (2×P4M)  -GFP-CIBN-HOTag6 | P4M: deep yellow GFP: green CIBN: black linker-HOTag6: blue  MTASTENFKNVKEKYQQMRGDALKTEILADFKDKLAEATDEQSLKQIVAELKSKDEYRILAKGQGLTTQLLGLKTSSVSSFEKMVEETRESIKSQERQTIKIKGGMVSKGEELFTGVVPILVELDGDVNGHKFSVSGEGEGDATYGKLTLKFICTTGKLPVPWPTLVTTLTYGVQCFARYPDHMKQHDFFKSAMPEGYVQERTIFFKDDGNYKTRAEVKFEGDTLVNRIELKGIDFKEDGNILGHKLEYNYNSHKVYITADKQKNGIKVNFKTRHNIEDGSVQLADHYQQNTPIGDGPVLLPDNHYLSTQSKLSKDPNEKRDHMVLLEFVTAAGITLGMDELYASGSGSMNGAIGGDLLLNFPDMSVLERQRAHLKYLNPTFDSPLAGFFADSSMITGGEMDSYLSTAGLNLPMMYGETTVEGDSRLSISPETTLGTGNFKAAKFDTETKDCNEAAKKMTMNRDDLVEEGEEEKSKITEQNNGSTKSIKKMKHKAKKEENNFSNDSSKVTKELEKTDYIHVGSGSAGGSAGGSAGGSAGGSAGGSAGGSAGGSRTLREIEELLRKIIEDSVRSVAELEDIEKWLKKI  **2×P4M:**  MTASTENFKNVKEKYQQMRGDALKTEILADFKDKLAEATDEQSLKQIVAELKSKDEYRILAKGQGLTTQLLGLKTSSVSSFEKMVEETRESIKSQERQTIKIKGGGSGGGSTASTENFKNVKEKYQQMRGDALKTEILADFKDKLAEATDEQSLKQIVAELKSKDEYRILAKGQGLTTQLLGLKTSSVSSFEKMVEETRESIKSQERQTIKIK |
| pLenti-mCh-CRY2-  HOTag3 | mCherry:deep red CRY2:deep blue linker:red linker-HOTag3: blue  MVSKGEEDNMAIIKEFMRFKVHMEGSVNGHEFEIEGEGEGRPYEGTQTAKLKVTKGGPLPFAWDILSPQFMYGSKAYVKHPADIPDYLKLSFPEGFKWERVMNFEDGGVVTVTQDSSLQDGEFIYKVKLRGTNFPSDGPVMQKKTMGWEASSERMYPEDGALKGEIKQRLKLKDGGHYDAEVKTTYKAKKPVQLPGAYNVNIKLDITSHNEDYTIVEQYERAEGRHSTGGMDELYKRSRSAAAGAGGAARAMKMDKKTIVWFRRDLRIEDNPALAAAAHEGSVFPVFIWCPEEEGQFYPGRASRWWMKQSLAHLSQSLKALGSDLTLIKTHNTISAILDCIRVTGATKVVFNHLYDPVSLVRDHTVKEKLVERGISVQSYNGDLLYEPWEIYCEKGKPFTSFNSYWKKCLDMSIESVMLPPPWRLMPITAAAEAIWACSIEELGLENEAEKPSNALLTRAWSPGWSNADKLLNEFIEKQLIDYAKNSKKVVGNSTSLLSPYLHFGEISVRHVFQCARMKQIIWARDKNSEGEESADLFLRGIGLREYSRYICFNFPFTHEQSLLSHLRFFPWDADVDKFKAWRQGRTGYPLVDAGMRELWATGWMHNRIRVIVSSFAVKFLLLPWKWGMKYFWDTLLDADLECDILGWQYISGSIPDGHELDRLDNPALQGAKYDPEGEYIRQWLPELARLPTEWIHHPWDAPLTVLKASGVELGTNYAKPIVDIDTARELLAKAISRTREAQIMIGAAARGAAAGAGGAGRGGGGSGSGSAGGSAGGSAGGSAGGSAGGSAGGSAGGSRGEIAKSLKEIAKSLKEIAWSLKEIAKSLKG |
| pLenti-2×P4M-mGFP  -CIBN-HOTag6-P2A-CRY2-HOTag3 | P4M: deep yellow linker: red mGFP: green CIBN: black  linker-HOTag6: blue P2A: purple CRY2:deep blue  linker-HOTag3: yellow  MTASTENFKNVKEKYQQMRGDALKTEILADFKDKLAEATDEQSLKQIVAELKSKDEYRILAKGQGLTTQLLGLKTSSVSSFEKMVEETRESIKSQERQTIKIKGGGSGGGSTASTENFKNVKEKYQQMRGDALKTEILADFKDKLAEATDEQSLKQIVAELKSKDEYRILAKGQGLTTQLLGLKTSSVSSFEKMVEETRESIKSQERQTIKIKGPGSGSGSMVSKGEELFTGVVPILVELDGDVNGHKFSVSGEGEGDATYGKLTLKFICTTGKLPVPWPTLVTTLTYGVQCFARYPDHMKQHDFFKSAMPEGYVQERTIFFKDDGNYKTRAEVKFEGDTLVNRIELKGIDFKEDGNILGHKLEYNYNSHKVYITADKQKNGIKVNFKTRHNIEDGSVQLADHYQQNTPIGDGPVLLPDNHYLSTQSKLSKDPNEKRDHMVLLEFVTAAGITLGMDELYASGSGSMNGAIGGDLLLNFPDMSVLERQRAHLKYLNPTFDSPLAGFFADSSMITGGEMDSYLSTAGLNLPMMYGETTVEGDSRLSISPETTLGTGNFKAAKFDTETKDCNEAAKKMTMNRDDLVEEGEEEKSKITEQNNGSTKSIKKMKHKAKKEENNFSNDSSKVTKELEKTDYIHVGSGSAGGSAGGSAGGSAGGSAGGSAGGSAGGSRTLREIEELLRKIIEDSVRSVAELEDIEKWLKKIGSGATNFSLLKQAGDVEENPGPMKMDKKTIVWFRRDLRIEDNPALAAAAHEGSVFPVFIWCPEEEGQFYPGRASRWWMKQSLAHLSQSLKALGSDLTLIKTHNTISAILDCIRVTGATKVVFNHLYDPVSLVRDHTVKEKLVERGISVQSYNGDLLYEPWEIYCEKGKPFTSFNSYWKKCLDMSIESVMLPPPWRLMPITAAAEAIWACSIEELGLENEAEKPSNALLTRAWSPGWSNADKLLNEFIEKQLIDYAKNSKKVVGNSTSLLSPYLHFGEISVRHVFQCARMKQIIWARDKNSEGEESADLFLRGIGLREYSRYICFNFPFTHEQSLLSHLRFFPWDADVDKFKAWRQGRTGYPLVDAGMRELWATGWMHNRIRVIVSSFAVKFLLLPWKWGMKYFWDTLLDADLECDILGWQYISGSIPDGHELDRLDNPALQGAKYDPEGEYIRQWLPELARLPTEWIHHPWDAPLTVLKASGVELGTNYAKPIVDIDTARELLAKAISRTREAQIMIGAAARGAAAGAGGAGRGGGGSGSGSAGGSAGGSAGGSAGGSAGGSAGGSAGGSRGEIAKSLKEIAKSLKEIAWSLKEIAKSLKG |
| pLenti-NES-2×PABP-mGFP-CIBN-HOTag6-P2A-CRY2-HOTag3 | NES-2×PABP:deep yellow linker: red mGFP:green CIBN: black  linker-HOTag6: blue P2A: purple CRY2:deep blue  linker-HOTag3: yellow  MSRANSNELALKLAGLDINKTESRMDNCSASRRRDRLHVKLKSLRNKIHKQLHPNCRFDDATKTSGGGSGGGSMDNCSGSRRRDRLHVKLKSLRNKIHKQLHPNCRFDDATKTSGPGSGSGSMVSKGEELFTGVVPILVELDGDVNGHKFSVSGEGEGDATYGKLTLKFICTTGKLPVPWPTLVTTLTYGVQCFARYPDHMKQHDFFKSAMPEGYVQERTIFFKDDGNYKTRAEVKFEGDTLVNRIELKGIDFKEDGNILGHKLEYNYNSHKVYITADKQKNGIKVNFKTRHNIEDGSVQLADHYQQNTPIGDGPVLLPDNHYLSTQSKLSKDPNEKRDHMVLLEFVTAAGITLGMDELYASGSGSMNGAIGGDLLLNFPDMSVLERQRAHLKYLNPTFDSPLAGFFADSSMITGGEMDSYLSTAGLNLPMMYGETTVEGDSRLSISPETTLGTGNFKAAKFDTETKDCNEAAKKMTMNRDDLVEEGEEEKSKITEQNNGSTKSIKKMKHKAKKEENNFSNDSSKVTKELEKTDYIHVGSGSAGGSAGGSAGGSAGGSAGGSAGGSAGGSRTLREIEELLRKIIEDSVRSVAELEDIEKWLKKIGSGATNFSLLKQAGDVEENPGPMKMDKKTIVWFRRDLRIEDNPALAAAAHEGSVFPVFIWCPEEEGQFYPGRASRWWMKQSLAHLSQSLKALGSDLTLIKTHNTISAILDCIRVTGATKVVFNHLYDPVSLVRDHTVKEKLVERGISVQSYNGDLLYEPWEIYCEKGKPFTSFNSYWKKCLDMSIESVMLPPPWRLMPITAAAEAIWACSIEELGLENEAEKPSNALLTRAWSPGWSNADKLLNEFIEKQLIDYAKNSKKVVGNSTSLLSPYLHFGEISVRHVFQCARMKQIIWARDKNSEGEESADLFLRGIGLREYSRYICFNFPFTHEQSLLSHLRFFPWDADVDKFKAWRQGRTGYPLVDAGMRELWATGWMHNRIRVIVSSFAVKFLLLPWKWGMKYFWDTLLDADLECDILGWQYISGSIPDGHELDRLDNPALQGAKYDPEGEYIRQWLPELARLPTEWIHHPWDAPLTVLKASGVELGTNYAKPIVDIDTARELLAKAISRTREAQIMIGAAARGAAAGAGGAGRGGGGSGSGSAGGSAGGSAGGSAGGSAGGSAGGSAGGSRGEIAKSLKEIAKSLKEIAWSLKEIAKSLKG |
| pLenti-CMV-cpLOV2-NLS-mGFP-NES2(L  5A+I14A)-2×PABD(Opi1p) | cpLOV2:yellow  linker:red NES1:gray mGFP:green  NES2(L5A and I14A):gray Opi1p:deep yellow NLS:blue  MTEHVRDAAEREGVMLIKKTAENIDEAAKELAGLDLGGGSGGSGGGLATTLERIEKNFVITDPRLPDNPIIFASDSFLQLTEYSREEILGRNCRFLQGPETDRATVRKIRDAIDNQTEVTVQLINYTKSGKKFWNLFHLQPMRDQKGDVQYFIGVQLDGGSGSGSAAAKRSWSMAFGSGSGSMVSKGEELFTGVVPILVELDGDVNGHKFSVSGEGEGDATYGKLTLKFICTTGKLPVPWPTLVTTLTYGVQCFSRYPDHMKQHDFFKSAMPEGYVERTIFFKDDGNYKTRAEVKFEGDTLVNRIELKGIDFKEDGNILGHKLEYNYNSHNVYIMADKQKNGIKVNFKIRHNIEDGSVQLADHYQQNTPIGDGPVLLPDNHYLSTQSKLSKDPNEKRDHMVLLEFVTAAGITLGMDELYKSGLRSRANSNE**A**ALKLAGLD**A**NKTESRGSGSGSKRQKLSRAIAKGKDNLKEYKLNMSIESKKRLVTCLHLLKLANKQLSDKISCLQDLVEKEQGSGSKRQKLSRAIAKGKDNLKEYKLNMSIESKKRLVTCLHLLKLANKQLSDKISCLQDLVEKEQ |
| pLenti-CMV-cpLOV2-NLS-mGFP-NES2(L5A+I14A)-3×PABP(Raf1) | cpLOV2:yellow linker:red NES1:gray mGFP:green  NES2(L5A and I14A):gray Raf1:deep yellow NLS:blue  MTEHVRDAAEREGVMLIKKTAENIDEAAKELAGLDLGGGSGGSGGGLATTLERIEKNFVITDPRLPDNPIIFASDSFLQLTEYSREEILGRNCRFLQGPETDRATVRKIRDAIDNQTEVTVQLINYTKSGKKFWNLFHLQPMRDQKGDVQYFIGVQLDGGSGSGSAAAKRSWSMAFGSGSGSMVSKGEELFTGVVPILVELDGDVNGHKFSVSGEGEGDATYGKLTLKFICTTGKLPVPWPTLVTTLTYGVQCFSRYPDHMKQHDFFKSAMPEGYVQERTIFFKDDGNYKTRAEVKFEGDTLVNRIELKGIDFKEDGNILGHKLEYNYNSHNVYIMADKQKNGIKVNFKIRHNIEDGSVQLADHYQQNTPIGDGPVLLPDNHYLSTQSKLSKDPNEKRDHMVLLEFVTAAGITLGMDELYKSGLRSRANSNE**A**ALKLAGLD**A**NKTESRPEQFQAFRNEVAVLRKTRHVNILLFMGYMTKDNLAIVTQWCEGGGGSGGGSPEQFQAFRNEVAVLRKTRHVNILLFMGYMTKDNLAIVTQWCEGGGGSGGGSPEQFQAFRNEVAVLRKTRHVNILLFMGYMTKDNLAIVTQWCEG |
| pLenti-CMV-cpLOV2  -NLS-mGFP-NES2(L5A+I14A)-2×PABP(4E mutations) | cpLOV2:yellow linker:red NES1:gray mCherry:shiny red  NES2(WT):gray PABD(4E mutation):deep yellow NLS:blue  MTEHVRDAAEREGVMLIKKTAENIDEAAKELAGLDLGGGSGGSGGGLATTLERIEKNFVITDPRLPDNPIIFASDSFLQLTEYSREEILGRNCRFLQGPETDRATVRKIRDAIDNQTEVTVQLINYTKSGKKFWNLFHLQPMRDQKGDVQYFIGVQLDGGSGSGSAAAKRSWSMAFGSGSGSMVSKGEEDNMAIIKEFMRFKVHMEGSVNGHEFEIEGEGEGRPYEGTQTAKLKVTKGGPLPFAWDILSPQFMYGSKAYVKHPADIPDYLKLSFPEGFKWERVMNFEDGGVVTVTQDSSLQDGEFIYKVKLRGTNFPSDGPVMQKKTMGWEASSERMYPEDGALKGEIKQRLKLKDGGHYDAEVKTTYKAKKPVQLPGAYNVNIKLDITSHNEDYTIVEQYERAEGRHSTGGMDELYKSGLRSRANSNE**A**ALKLAGLD**A**NKTESRMDNCSASRRRDRLHVELESLENEIHKQLHPNCRFDDATKTSGGGSGGGSMDNCSASRRRDRLHVELESLENEIHKQLHPNCRFDDATKTSSWWWWZW |
| pLenti-CMV-  mCherry-PLD2 | mCherry: shiny red linker: red PLD2: blue  MVSKGEEDNMAIIKEFMRFKVHMEGSVNGHEFEIEGEGEGRPYEGTQTAKLKVTKGGPLPFAWDILSPQFMYGSKAYVKHPADIPDYLKLSFPEGFKWERVMNFEDGGVVTVTQDSSLQDGEFIYKVKLRGTNFPSDGPVMQKKTMGWEASSERMYPEDGALKGEIKQRLKLKDGGHYDAEVKTTYKAKKPVQLPGAYNVNIKLDITSHNEDYTIVEQYERAEGRHSTGGMDELYKGGGSGGGSGSMTATPESLFPTGDELDSSQLQMESDEVDTLKEGEDPADRMHPFLAIYELQSLKVHPLVFAPGVPVTAQVVGTERYTSGSKVGTCTLYSVRLTHGDFSWTTKKKYRHFQELHRDLLRHKVLMSLLPLARFAVAYSPARDAGNREMPSLPRAGPEGSTRHAASKQKYLENYLNRLLTMSFYRNYHAMTEFLEVSQLSFIPDLGRKGLEGMIRKRSGGHRVPGLTCCGRDQVCYRWSKRWLVVKDSFLLYMCLETGAISFVQLFDPGFEVQVGKRSTEARHGVRIDTSHRSLILKCSSYRQARWWAQEITELAQGPGRDFLQLHRHDSYAPPRPGTLARWFVNGAGYFAAVADAILRAQEEIFITDWWLSPEVYLKRPAHSDDWRLDIMLKRKAEEGVRVSILLFKEVELALGINSGYSKRALMLLHPNIKVMRHPDQVTLWAHHEKLLVVDQVVAFLGGLDLAYGRWDDLHYRLTDLGDSSESAASQPPTPRPDSPATPDLSHNQFFWLGKDYSNLITKDWVQLDRPFEDFIDRETTPRMPWRDVGVVVHGLPARDLARHFIQRWNFTKTTKAKYKTPTYPYLLPKSTSTANQLPFTLPGGQCTTVQVLRSVDRWSAGTLENSILNAYLHTIRESQHFLYIENQFFISCSDGRTVLNKVGDEIVDRILKAHKQGWCYRVYVLLPLLPGFEGDISTGGGNSIQAILHFTYRTLCRGEYSILHRLKAAMGTAWRDYISICGLRTHGELGGHPVSELIYIHSKVLIADDRTVIIGSANINDRSLLGKRDSELAVLIEDTETEPSLMNGAEYQAGRFALSLRKHCFGVILGANTRPDLDLRDPICDDFFQLWQDMAESNANIYEQIFRCLPSNATRSLRTLREYVAVEPLATVSPPLARSELTQVQGHLVHFPLKFLEDESLLPPLGSKEGMIPLEVWT |
| pLVX-TR3GS-Tom20-mGFP (or none)-PLD  _PMF100×_-hPGK-TetOne | Tom20:purple mGFP:green PLD_PMF100×_:orange  linker:red  MVGRNSAIAAGVCGALFIGYCIYFDRKRRSDPNFKNRLRERRKKQKLAKERAGLSKLPDLKDAEAVQKFFLEEIQLGEELLAQGEYEKGVDHLTNAIAVCGQPQQLLQVLQQTLPPPVFQMLLTKLPTISQRIVSAQSLAEDDVEGGSGDPPVATMVSKGEELFTGVVPILVELDGDVNGHKFSVSGEGEGDATYGKLTLKFICTTGKLPVPWPTLVTTLTYGVQCFARYPDHMKQHDFFKSAMPEGYVQERTIFFKDDGNYKTRAEVKFEGDTLVNRIELKGIDFKEDGNILGHKLEYNYNSHKVYITADKQKNGIKVNFKTRHNIEDGSVQLADHYQQNTPIGDGPVLLPDNHYLSTQSKLSKDPNEKRDHMVLLEFVTAAGITLGMDELYARGAAAGAGGAGRGGGGSADSATPHLDAVEQTLRQVSPGLEGDVWERTSGNKLDGSAADPSDWLLQTPGCWGDDRCVDRVGTKRLLAKMTENIGNATRTVDISTLAPFPNGAFQDAIVAGLKESAARGNKLKVRILVGAAPVYHMNVIPSKYRDELTAKLGKAAENITLNVASMTTSKTAFSWNHSKILVVDGQSALTGGINSWKDDYLDTTHPVSDVDLALTGPAAGSAGRYLDTLWTWTCQNKSNIASVWFAASGNAGCMATMHKDTNPKASPATGNVPIIAVGGLGVGIKDVDPKSTFRPDLPTASDTKCVVGLHDNTNADRDYDTVNPEESALRALVASAKSHIEISQQDLNATCPPLPRYDIRLYDALAAKMAAGVKVRIVVSDPANRGAVGSVGYSQIKSLSEISDTLRNRLANITGSQQAAKTAMCSNLQLATFRSSPNDKWADGHPYAQHHKLVSVDSSTFYIGSKNLYPSWLQDFGYIVESPEAAKQLDAKLLDPQWKYSQETATVDYARGICNAEF |
